# Myosin-I Synergizes with Arp2/3 Complex to Enhance Pushing Forces of Branched Actin Networks

**DOI:** 10.1101/2024.02.09.579714

**Authors:** Mengqi Xu, David M. Rutkowski, Grzegorz Rebowski, Malgorzata Boczkowska, Luther W. Pollard, Roberto Dominguez, Dimitrios Vavylonis, E. Michael Ostap

## Abstract

Myosin-Is colocalize with Arp2/3 complex-nucleated actin networks at sites of membrane protrusion and invagination, but the mechanisms by which myosin-I motor activity coordinates with branched actin assembly to generate force are unknown. We mimicked the interplay of these proteins using the “comet tail” bead motility assay, where branched actin networks are nucleated by Arp2/3 complex on the surface of beads coated with myosin-I and the WCA domain of N-WASP. We observed that myosin-I increased bead movement efficiency by thinning actin networks without affecting growth rates. Remarkably, myosin-I triggered symmetry breaking and comet-tail formation in dense networks resistant to spontaneous fracturing. Even with arrested actin assembly, myosin-I alone could break the network. Computational modeling recapitulated these observations suggesting myosin-I acts as a repulsive force shaping the network’s architecture and boosting its force-generating capacity. We propose that myosin-I leverages its power stroke to amplify the forces generated by Arp2/3 complex-nucleated actin networks.

## Introduction

Actin assembly stimulated by Arp2/3 complex provides pushing forces for diverse cellular processes (*1–8*), including lamellipodial protrusion, endocytosis, phagocytosis, and cell adhesion. After being activated by membrane-associated nucleation promoting factors (NPFs), Arp2/3 complex nucleates new actin branches from the sides of pre-existing mother filaments, generating branched, dendritic actin networks that exert pushing forces against or deform the membrane. The network geometry, assembly kinetics, and mechanical properties are dynamically adapted by a set of actin binding proteins (e.g., capping proteins, nucleation promoting factors, profilin, cofilin, and crosslinkers) that respond to mechanical loading (*9–19*).

Class-I myosins (myosin-Is) (*20, 21*) frequently colocalize with Arp2/3 complex-nucleated branched actin networks near membranes (*22–26*). As an actin-activated ATPase, the myosin-I motor dynamically detaches and attaches from actin filaments in an ATP-dependent manner while generating force through a lever arm-mediated power stroke (*27*). Myosin-Is are single-headed, membrane-anchored motors that bind directly to cell membranes and function in membrane deformation, cell adhesion, and intracellular trafficking (*21*). Although multiple cellular studies have shown that the myosin-I motor activity cooperates with dynamic actin assembly by Arp2/3 complex at the plasma membrane, the functional outcome of this interaction remains unknown.

To investigate the functional interaction between Arp2/3 complex and myosin-I in actin assembly, we developed a biomimetic system using a comet-tail bead motility assay (*9, 12, 28*). This system recapitulates the colocalization of NPFs and myosin-I observed on cellular membranes by employing micron-sized beads coated with both NPF and myosin-I. We investigated the impact of myosin-I on Arp2/3 complex-mediated branched actin assembly at varying network densities achieved through different capping protein (CP) concentrations (*12, 14, 29*).

Our findings revealed that myosin-I alters actin assembly kinetics by reducing Arp2/3 complex-mediated branching at NPF-coated surfaces, resulting in a sparser actin network that exhibits enhanced elongation efficiency. This effect is attributed to the pushing force exerted by the myosin-I power stroke directly on surrounding actin networks, propelling actin filaments away. Furthermore, the myosin-I power stroke generates sufficient force to disrupt the network and trigger actin shell breakdown. Notably, a computational model at the molecular level provides mechanistic insights, suggesting that myosin-I promotes force generation during Arp2/3 complex-mediated actin polymerization by exerting a repulsive force on the branched network via its power stroke.

## Results

### Comet-tail bead motility asshay

The comet-tail bead motility assay was used to investigate the effect of myosin-I activity on Arp2/3 complex-mediated branched actin assembly (*9, 28*). Full-length, biotinylated *Drosophila* Myo1d was attached to the bead surface via neutravidin (myosin-bead; see Methods; Fig. 1A) alongside a GST-tagged WCA domain from human N-WASP. *Drosophila* Myo1d was chosen as it functions optimally at 20-22 °C (*30, 31*). Bead-bound Myo1d activity was confirmed by processive bead movement along actin filaments (Fig. 1B; Movie S1). Control-beads were made identically to the myosin-beads, except that a biotinylated far-red fluorescent dye (CF640, see Methods) was bound to the neutravidin (Fig. 1A). Actin assembly around myosin- and control-beads was imaged simultaneously by epifluorescence microscopy (Fig. 1C).

**Figure 1.**
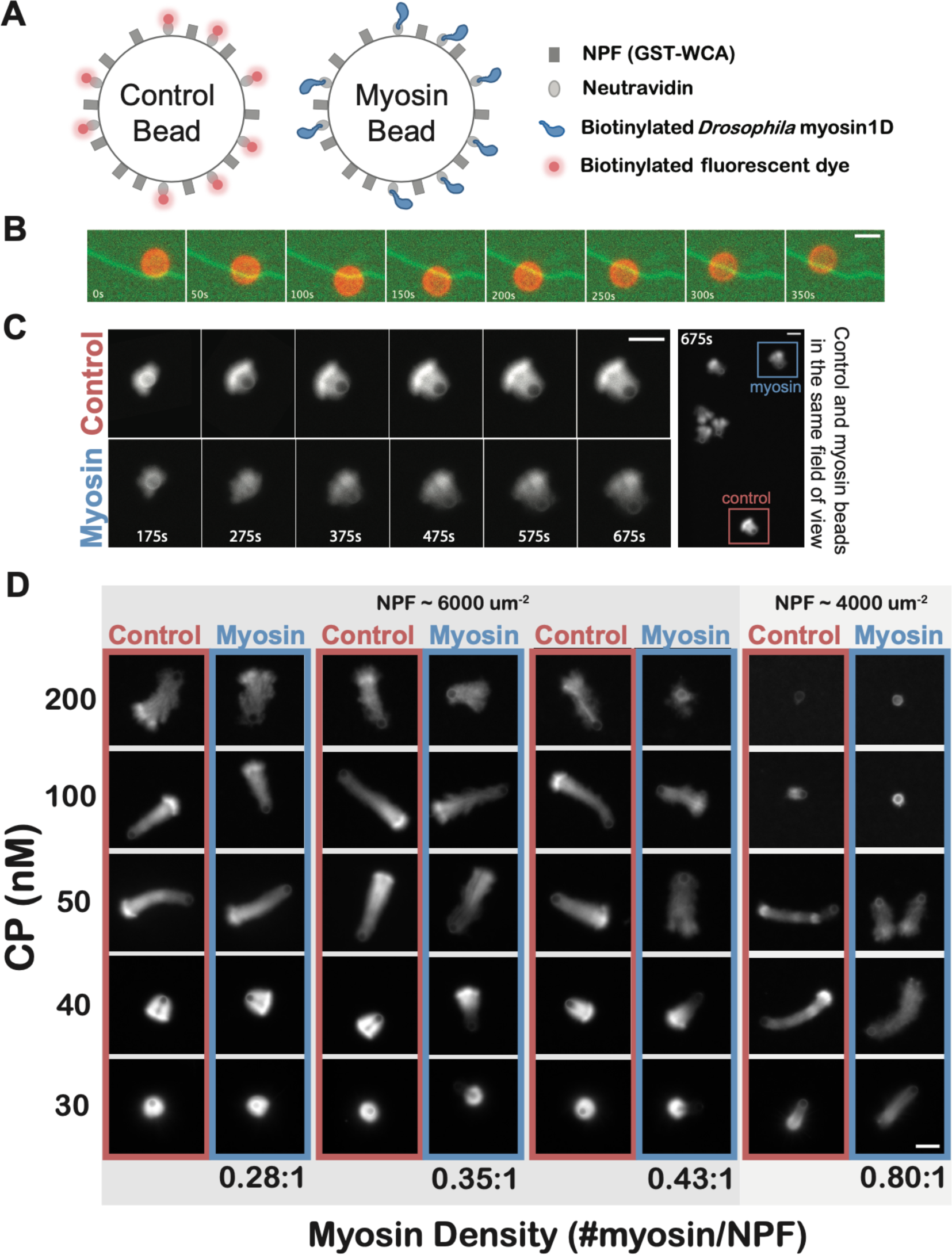
Alteration of comet tail morphology by myosin-I. **(A)** Schematic of (left) control- and (right) myosin-beads coated with NPF and neutravidin, where neutravidin is further conjugated with (control-bead) biotinylated CF640 fluorescent dye or (myosin-bead) biotinylated *Drosophila* Myo1d. (**B**) Time-lapse sequence of a (red) myosin-bead walking on a (green) single-actin-filament track. Speed ∼8 nm/s. Scale bar 2 µm. (**C**) Time series of the growth of actin comet tail from symmetry breaking to generation of a polarized comet tail from (top) control-bead and (bottom) myosin-bead. Scale bar 5 µm. (**D**) Phase map showing representative examples of 2 µm-diameter beads coated with a range of myosin-densities in the presence of 30 – 200 nM CP, 4 µM (5% Rhodamine labeled) actin, and 200 nM Arp2/3 complex. For each myosin density condition, the left column show control-beads that were acquired in the same imaging field as the myosin-beads in the right column. The myosin and NPF densities were determined by SDS-PAGE gel, where the NPF density is the same for group 0.28:1, 0.35:1 and 0.43:1, ∼ 6000 um^−2^, and ∼ 4000 um^−2^ for group 0.80:1. Images were acquired 20-35 min after mixing. Brightness and contrast were linearly set to be the same for each control- and myosin-bead pair but different among panels at different conditions for better visualization. Scale bar 5 µm.

Upon mixing of assay components, surface-bound NPFs stimulate Arp2/3 complex-mediated branched actin assembly. Polymerization at the bead surface displaces the branched actin arrays outwards, building tension in the actin network, and ultimately breaking symmetry and forming polarized comet tails (*9, 14, 32–35*) (Fig. 1C). By varying CP concentration, we created networks of varying actin density that ranged from tightly packed and symmetry-breaking resistant (dense) to loosely woven and fracture-prone (sparse), allowing us to investigate Myo1d’s influence on symmetry breaking, comet tail formation, and morphology.

The morphology of comet tails generated by myosin-beads differed from control-beads at all network densities achieved by varying CP concentrations (Fig. 1D; Fig. S1), and these differences were more pronounced at higher Myo1d densities. We identified three distinct regimes: high, intermediate, and low network densities, controlled by the CP concentration (200 nM, 40 nM-100 nM, and 30 nM, respectively) (Fig. 1D; Fig. S1), and describe each regime in greater detail using the 0.43 myosin / 1 NPF ratio in the following sections.

### Myosin-I prevents comet tail formation from sparse actin networks

Under high CP concentrations (200 nM), actin elongation was suppressed by capping, resulting in actin arrays consisting of short branches. As a result, actin comet tails grown from control-beads were irregularly organized with sparse networks (Fig. 1D; Fig. 2A), consistent with previous reports (*12, 14, 36*). Myosin-beads (0.43:1 myosin/NPF) failed to generate a comet tail (Fig. 2A; Movie S2), but instead formed loose actin clouds which diffused away from the bead when flow was present in the chamber. The presence of myosin compromised the network cohesion, making it more prone to disruption, and inhibiting comet tail formation.

**Figure 2.**
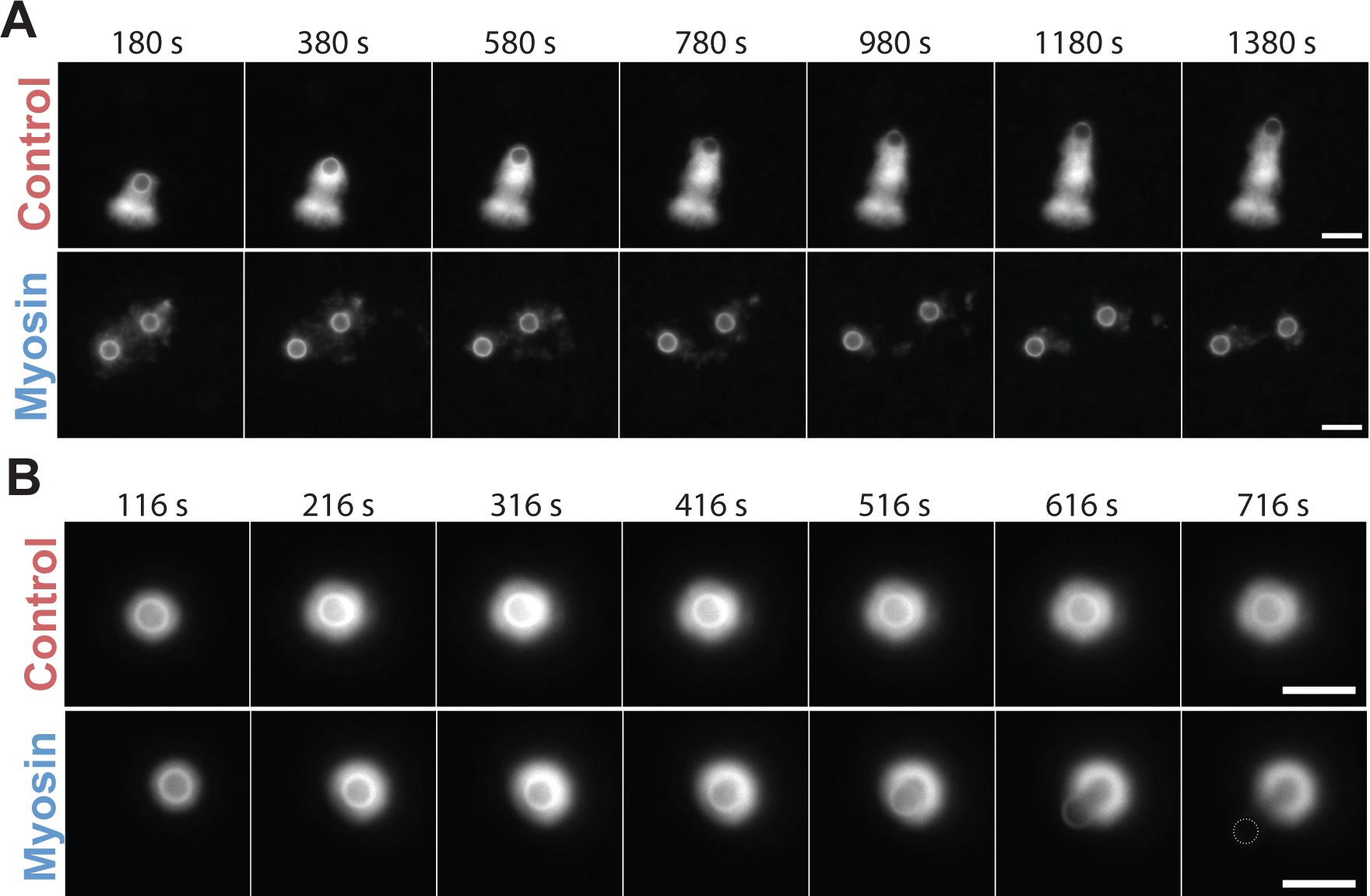
Disruption and fracturing of actin shells by myosin-I. (**A**) Time series of (top) control-beads and (bottom) 0.43:1 myosin-beads acquired in the same imaging field in the presence of 200 nM CP showing the inability of myosin-beads to form a comet tail. (**B**) Time series of (top) control-beads and (bottom) 0.43:1 myosin-beads acquired in the same field in the presence of 25 nM CP showing fracturing of the actin shell and comet tail growth from a myosin-bead but not the control-bead. Conditions: 4 µM actin (5% Rhodamine labeled), 200 nM Arp2/3 complex, 200 nM or 25 nM CP. Scale bar 5 µm.

### Myosin-I facilitates fracturing of dense actin networks

At a low CP concentration (25 nM), long actin filaments emanating from bead surfaces become entangled, forming dense actin networks that are highly resistant to fracture by actin polymerization forces (*12, 14, 34, 36*). Control-beads remained encapsulated within the shell, without symmetry breaking, for > 15 minutes after the initiation of the polymerization (Fig. 1D; Fig. 2B). Strikingly, myosin-beads grown under the same condition broke symmetry within 10 min of mixing (Fig. 2B; Movie S3). This result suggests that myosin-I may enhance the force generation during actin assembly and/or may alter actin architecture that promotes network fracture and symmetry breaking.

### Myosin-I induces efficient comet tail growth

At intermediate CP concentrations (40 nM - 100 nM), myosin-beads generated comet tails with sparser actin networks compared to control-beads (Fig. 1D; Fig. S1; Fig. 3A-B; Movie. S4). Following the time course of tail elongation, we found that the sparser network architecture of myosin-beads arises from a 0.76-fold lower actin assembly rate (p <0.0001, n = 11, Fig. 3E-F) as quantified by measuring the rhodamine-actin fluorescence in the comet tail over time. Remarkably, despite the lower actin assembly rate, myosin comet tails in the presence of 50 nM CP elongated (0.69 ± 0.10 μm/min) at the same speed as the control-beads (0.66 ± 0.20 μm/min; Fig. 3C-D). If we define the growth efficiency of actin comet tail as the comet tail length divided by the total amount of actin incorporated, the myosin-beads exhibited a 1.4-fold higher growth efficiency than the control-beads (Fig. 3G).

**Figure 3.**
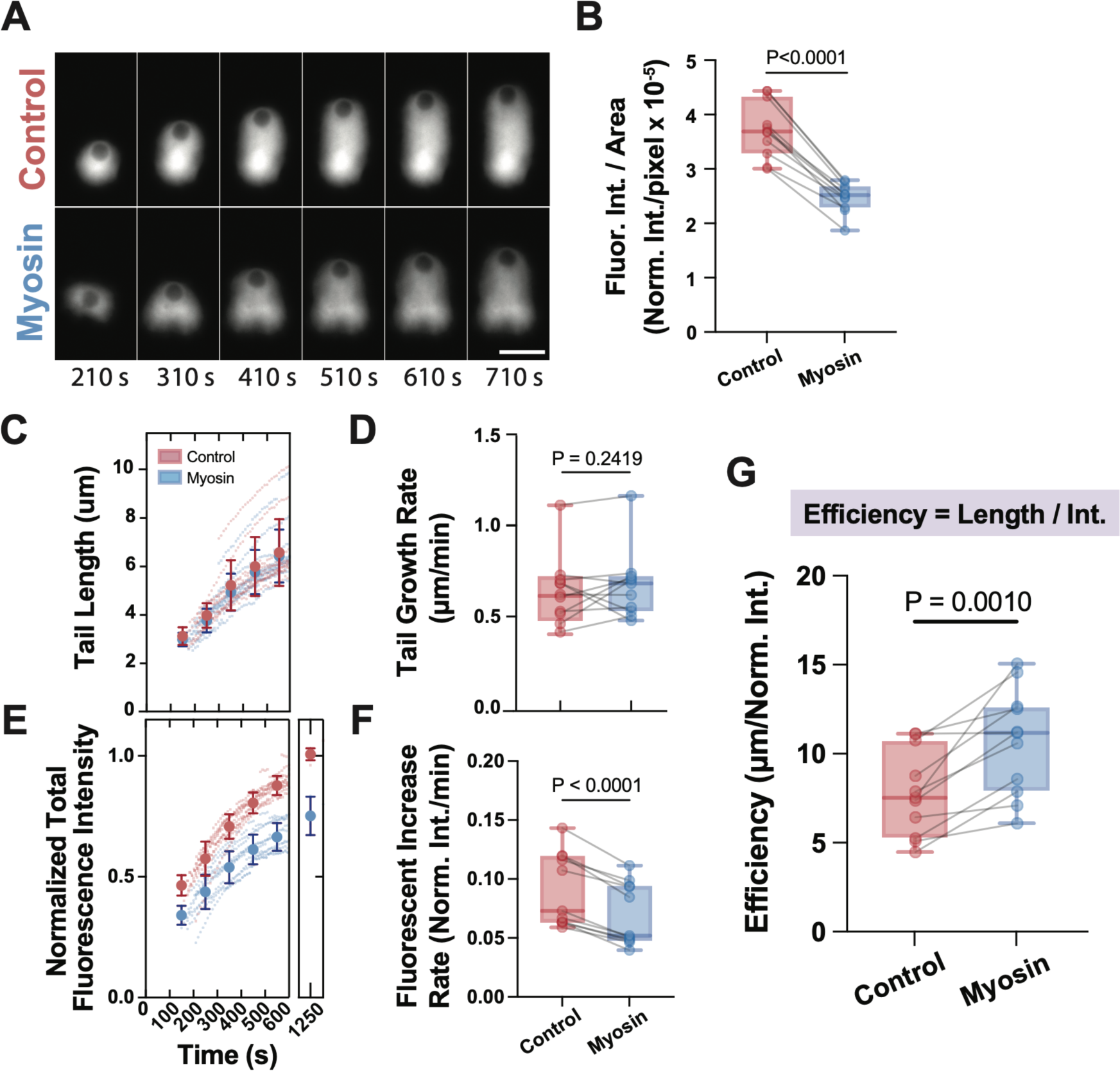
Myosin-I decreases the actin density of comet tails. (**A**) Time-series of (top) control- and (bottom) 0.43:1 myosin-beads growing comet tails in the presence of 50 nM CP. The actin network growing from the myosin-bead is less dense but has a similar tail length as the control. Conditions: 4 µM actin (5% Rhodamine labeled), 200 nM Arp2/3 complex and 50 nM CP. Scale bar 5 µm. **(B**) Comparison of the mean fluorescence intensities (Fluor. Int. / Area) of actin comet tails grown from control- and myosin-beads, captured 20 min after mixing. The solid lines connect experimental pairs (N=5, n = 11). (**C**) Actin comet tail length as a function of time and (**D**) tail growth rates derived from the slopes of the time courses in (**C**). (**E**) Comet tail fluorescence as a function of time. Control- and myosin-bead experimental pairs are normalized to the average fluorescence level of the control-beads from 1100 s –1300 s. (**F**) Rate of fluorescent actin incorporation into comet tails derived from the slopes of the time courses in (**E**). (**C, E**) Large points show the averaged value at binned time interval (every 100 sec). Traces are from individual beads with each myosin-bead acquired with a control-bead in the same field of view (N=5, n=11). Error bars are SD. (**G**) Growth efficiency for control and myosin-beads. Efficiency is defined as comet tail length per unit actin fluorescence intensity. Box plots (**B,D, F, G**) show median (center line), interquartile range (box) and min-max values (whiskers). p values were calculated using two-tail paired t test. Each point represents a control- and myosin-bead pair acquired in same field of view. See also Supplementary Fig.

We quantified the amount of a fluorescently labeled Arp2/3 complex (SNAP-Arp2/3 complex; see Methods) in the comet tail (Fig. 4A-C). Myosin-beads showed significantly reduced levels of SNAP-Arp2/3 complex (p<0.0001, n=33, Fig. 4C) in the network. This reduced Arp2/3 complex level also resulted in higher actin to Arp2/3 complex ratios on myosin-beads (Fig. 4D), indicating a sparse network organization with fewer branch points but longer filaments (*15*). Despite the reduced amount of Arp2/3 complex in the comet tails, we observed equivalent SNAP-Arp2/3 complex fluorescence on the control- and myosin-bead surfaces (Fig. 4E), suggesting that myosin-I does not affect the loading of Arp2/3 complex onto the bead-bound NPF. Rather, Arp2/3 complex is recruited to NPF-coated beads and is ready for the arrival of a mother filament and G-actin to initiate branched nucleation. Taken together, myosin-beads have less dense network structure as a result of reduced Arp2/3 complex-stimulated actin branching.

**Figure 4.**
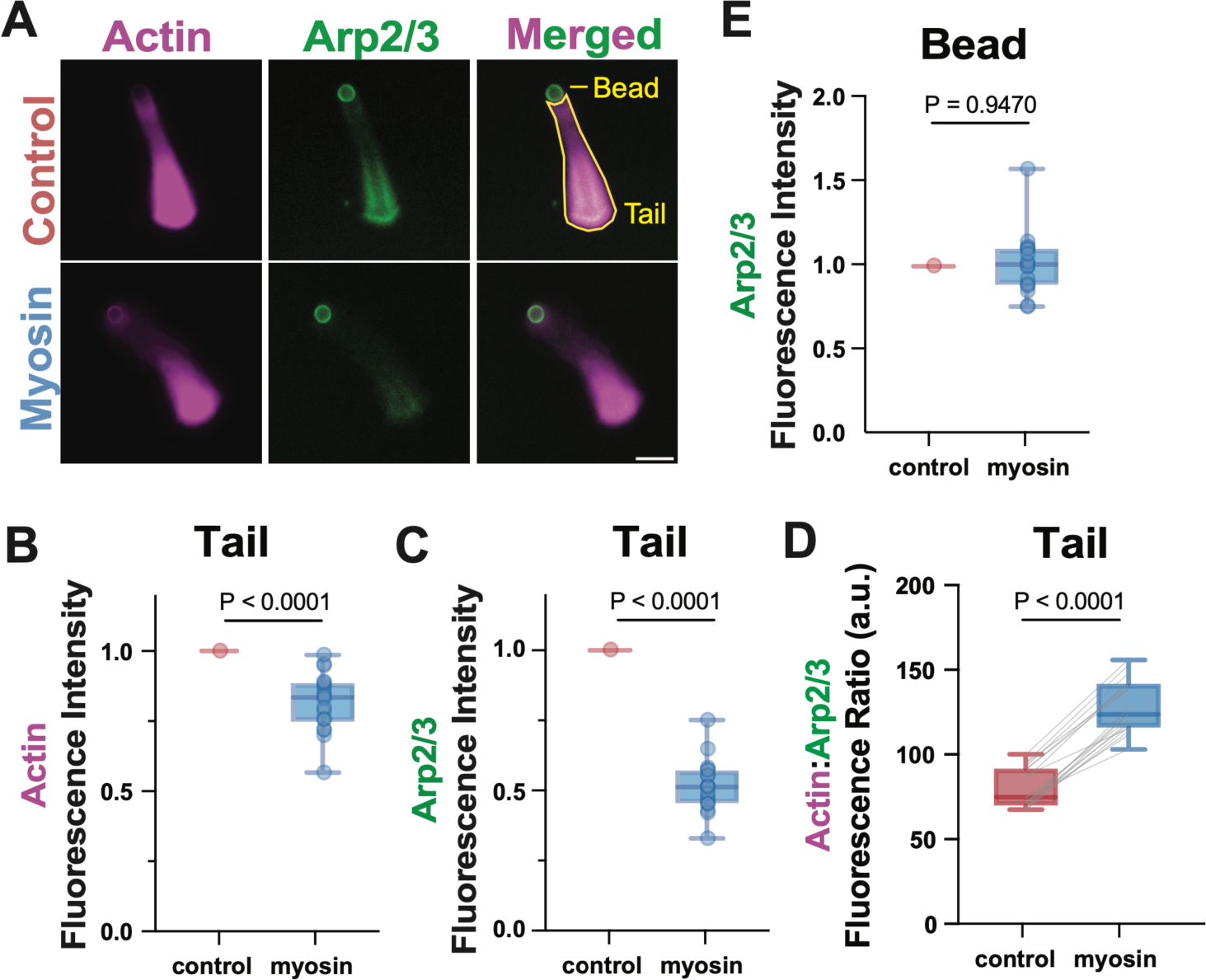
Comets grown from myosin-beads incorporate less Arp2/3 complex. (**A**) Representative actin comet tails assembled from (top) control- and (bottom) myosin-beads showing (magenta) actin and (green) SNAP-Arp2/3 complex. Images were acquired approximately 25 - 35 min after mixing. Brightness and contrast were set differently for actin and SNAP-Arp2/3 complex panels for visualization. Conditions: 4 µM actin (5% Rhodamine labeled), 200 nM Arp2/3 complex (80% SNAP-Surface 488 labeled), 40 nM CP. Scale bar is 5 µm. (**B**) Total actin fluorescence intensity, (**C**) total SNAP-Arp2/3 fluorescence intensity and (**D**) actin to SNAP-Arp2/3 fluorescence ratio over the entire comet tail region. (**E**) Total SNAP-Arp2/3 fluorescence intensity on bead surface. Plot shows median (center line), interquartile range (box) and min-max values (whiskers). p values were calculated using two-tail paired t test. Each point represents a pair of control- and myosin-beads (N=2, n=33)

We note that the myosin effect on growth efficiency depends on the CP concentration (Fig. 1D). Under low CP conditions (< 50 nM), myosin beads grew even longer tails than control-beads with sparse networks, suggesting an even higher growth efficiency than the 50 nM CP case as quantified above. High CP concentrations (> 50 nM) resulted in highly diffuse networks whose cohesion was easily compromised by myosin-I, causing actin dispersal, thus making their growth efficiency difficult to assess (*12, 14, 17*).

### The myosin power stroke is required to alter network architecture

We examined the role of myosin-I mechanochemical activity on actin assembly by removing ATP, resulting in the population of a long-lived, actin-bound, rigor state (i.e. rigor myosin) (*31*). To maintain normal actin polymerization, G-actin was pre-treated with ATP and gel filtered to eliminate free nucleotide (see Methods). Rigor myosin inhibited the formation of monopolar comet tails, while control-beads generated comet patterns as previously observed when free ATP was present (Fig. 5A-C). At 50 nM CP, rigor myosin-beads formed dense actin shells, which subsequently fractured, forming multiple short tails of high network density (Fig. 5A-C; Fig. S3C; Movie S5). Restoring myosin motor activity by adding ATP reproduced the sparse comet architecture with high growth efficiency as observed before (Fig. 5B; Fig. S3B). At the two extreme CP conditions (15 nM and 200 nM), rigor myosin-beads neither enhanced symmetry breaking nor shed actin away (Fig. 5C; Fig. S3D). Instead, actin shells were formed around the beads as a result of the strong actin-binding characteristics of rigor myosin. In addition to probing the effect of rigor myosin, we coupled biotinylated, Halo-tagged, actin-binding domain (Halo-ABD) of α-actinin to the beads, which binds actin filaments more dynamically than rigor myosin but does not undergo a power stroke (*31, 37, 38*). HaloABD-beads behaved similarly to rigor myosin (Fig. 5C; Fig. S3A; Movie. S6).

**Figure 5.**
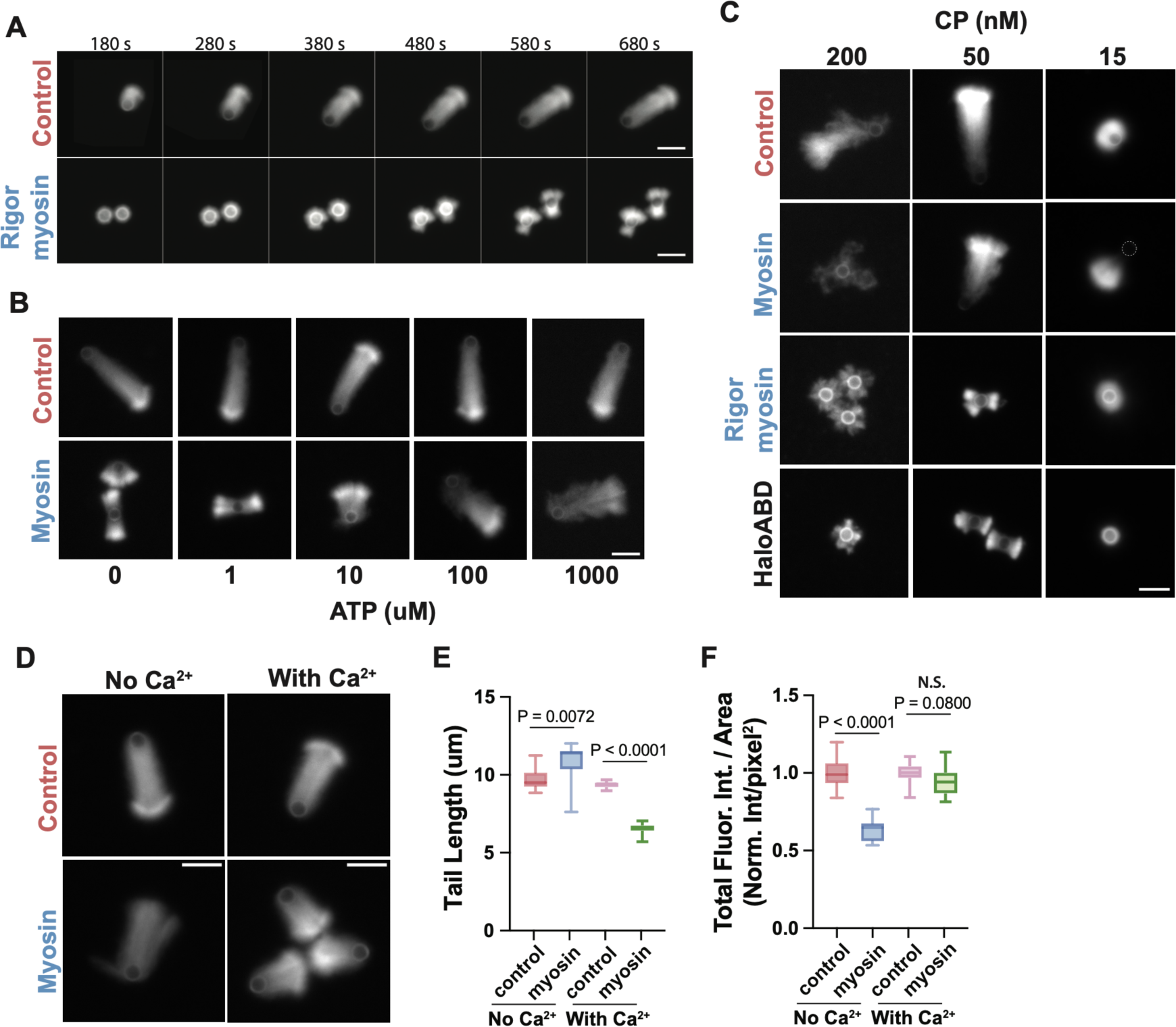
The myosin power stroke is required for altering network architectures. (**A**) Time-series of actin assembly around (top) control and (bottom) rigor myosin-beads at 50 nM CP. Rigor myosin heavily delayed the growth of actin comet tails. (**B**) Representative images of actin comet tails grown under different ATP concentrations (top: control beads; bottom: myosin-beads). Adding ATP back rescued comet tail growth. Images were acquired 20-30 min after mixing. (**C**) Representative actin network patterns assembled on control-, myosin-, rigor-myosin, and HaloABD-beads under three different CP concentrations, as indicated. Images were acquired 20 – 30 min after mixing. Brightness and contrast were set to the same values for each panel. (**D**) Representative actin comet tails generated by (top) control- and (bottom) myosin-beads in the absence and presence of 100 µM free Ca^2+^ at 50 nM CP. Images were acquired approximately 15-20 min after mixing. (**E**) Length and (**F**) network density quantified by total fluorescence intensity per area for control- and myosin-beads in the absence and presence of 100 µM free Ca^2+^ (N=1, n=15). Control- and myosin-bead experimental pairs are normalized to the average fluorescence level of the control-beads. Box plots (**E, F**) show median (center line), interquartile range (box) and min-max values (whiskers). p values were calculated using two-tail paired t test. Conditions: 4uM actin (5% Rhodamine labeled), 200 nM Arp2/3 complex, 15-200 nM CP, 0-1 mM ATP, as indicated. Scale bar 5 µm.

To further elucidate the role of the myosin mechanochemistry in modulating the actin network, we uncoupled the myosin actin-activated ATPase activity from the power stroke by adding calcium. Calcium weakens the affinity of the lever-arm stabilizing calmodulin light chains, resulting in an inhibited power stroke but preserved ATPase activity (*27, 39, 40*). We confirmed calcium inhibition using the *in vitro* gliding assay, where the Myo1d-powered F-actin gliding (speed 90.2±14.5 nm/s) was completely halted in the presence of 100 µM free calcium (Fig. S3E-F). Under these conditions, myosin-beads generated slightly shorter comet tails with network densities that were similar to control-beads (Fig. 5D-F). We conclude that myosin power stroke is required to alter the network architecture and induce a sparse actin organization with higher growth efficiency.

### The myosin power stroke alone can fracture the actin shell

We decoupled the force generated by myosin power stroke from the force generated by actin polymerization using Arp2/3 complex inhibitor, CK-666, and the polymerization inhibitor, Latrunculin B (LatB) (*12, 41*). The goal was to arrest actin assembly around the bead prior to the fracturing of the actin shell that results in symmetry breaking. Polymerization inhibitors were added ∼100 s after the initiation of polymerization. Strikingly, most of the myosin-beads fractured the actin shell and ejected the bead 15-20 min after polymerization arrest. In contrast, control-beads remained enclosed in the actin shell without observable changes during a 40 min time window (Fig. 6A; also see Fig. S4 and Movie S7-8 acquired in the absence of phalloidin).

**Figure 6.**
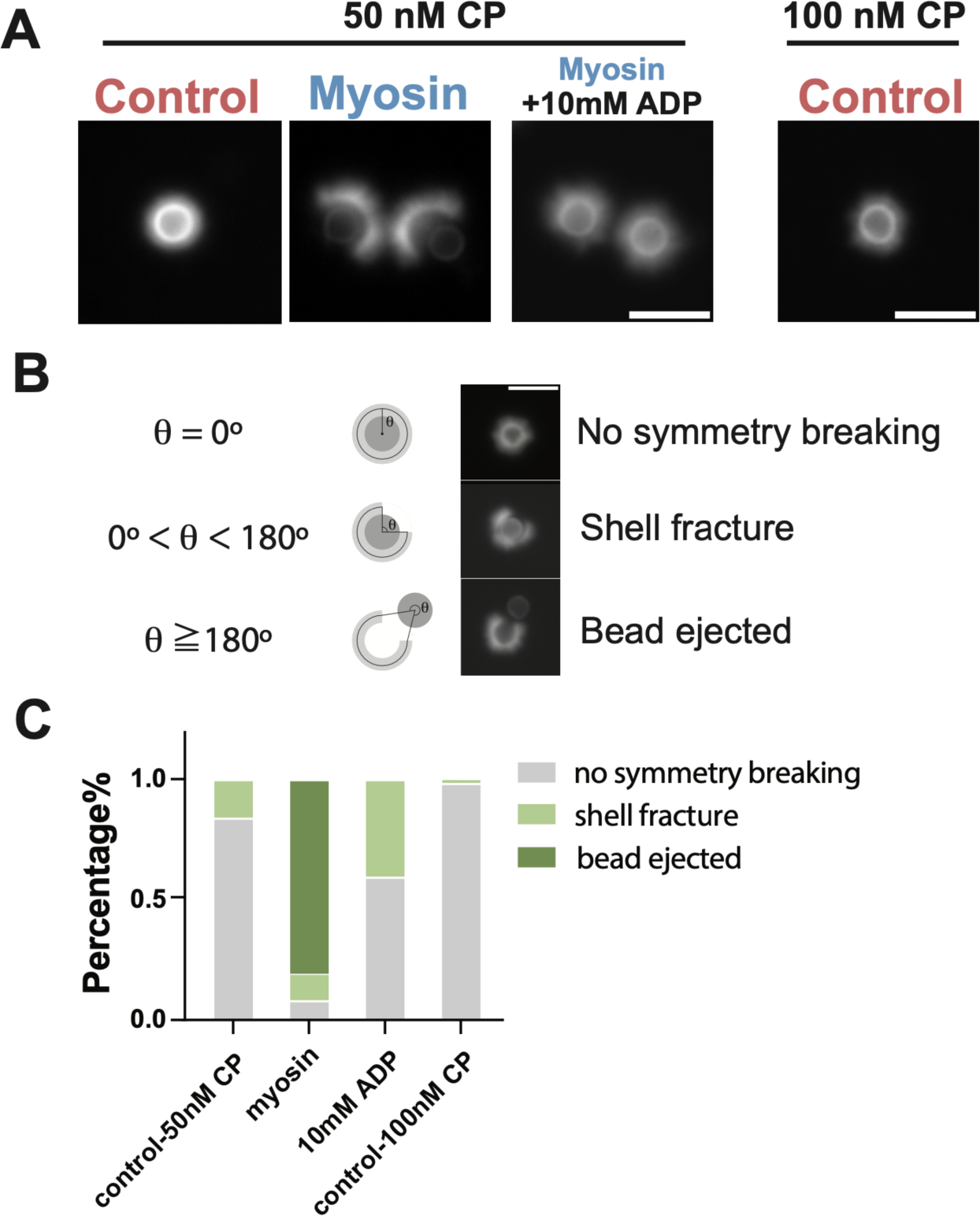
The myosin power stroke can fracture the actin shell. (**A**) Representative actin shells of control-, myosin-beads assembled under 50 nM CP, arrested by adding 20μM (5 molar excess) of Latrunculin B and CK-666 before symmetry breaking; as well as myosin-beads assemble under the same conditions but arrested with the addition of 10mM ADP to inhibit myosin power stroke. Myosin-beads fractured and ejected from the actin shell, while control- and myosin-beads (with 10mM ADP) remained enclosed in the shell. Control-bead assembled under 100 nM CP showing similar network density as the myosin-beads, didn’t show shell fracture or bead ejection. Image was captured approximately 40min after arrest. (**B**) The extents of shell breaking was classified by shell-breaking angle θ: θ = 0, no symmetry breaking; 0 < θ < 180, shell fracture; θ ≥ 180, bead ejected. Scale bar is 5μm. (**C**) Percentage of populations with different extents of shell breaking for control (50 nM CP) (n=182), myosin (n=158), myosin (with 10mM ADP) (n=89), control (100 nM CP) (n=65), N=2. Conditions: 4μM actin (5% Rhodamine labeled), 200 nM Arp2/3, 50 or 100 nM CP. 20μM (5 molar excess) phalloidin and 2 mM ATP were also added to prevent actin depolymerization and preserve myosin motor activity. Actin assembly was arrested 100s after mixing. Scale bar is 5 µm.

Shell fracture events were quantified by defining a shell-breaking angle, *θ*, where *θ* = 0, indicates no detected shell fracture; 0° < *θ* < 180°, indicates shell fracture; and *θ* ≥ 180° indicates the bead has been ejected from the shell (Fig. 6B). We found 80% of the myosin-beads were ejected from the shell and an additional 11% had fractured shells, while the corresponding control-beads showed only 16% shell fracture (Fig. 6C). Inhibition of myosin motile activity by adding 10 mM ADP (*31*) together with Latrunculin B and CK-666 resulted in substantial reduction in shell fracture events and eliminated bead ejection (Fig. 6A, C; Fig. S4C-D). Finally, we performed experiments with a higher CP concentration (100 nM) that results in control-beads of lower actin shell densities (Fig. 6A) and found only 2% shell fracture (Fig. 6C), which further confirmed that the shell breaking was a direct result of myosin power stroke, and is not attributed to differences in the shell network density.

### Simulations show myosin-I forces produce sparser actin networks and aid bead propulsion

We developed a filament-level computational model with an overall system size comparable to our experimental setup (Fig. 7A, see Methods, Table S1). We incorporated the myosin-I power stroke as a repulsive force that pushes actin away from the bead surface, and explore whether this myosin-induced pushing mechanism reproduces the experimentally observed comet tail patterns. In this model, we represented semiflexible actin filaments as beads connected by springs, polymerizing at their barbed ends and pushing against a spherical bead according to Brownian-ratchet-type force-elongation relationship. Spontaneous filament nucleation and branching at 70° angles occurs close to the bead; elongation stops by capping when filaments reach a specified length. The effect of fluid and bead-filament friction was combined into a single bead friction parameter. Excluded volume interactions prevent filament crossing, resulting in tensile and compressive stresses developing within a shell of branched actin filaments that nucleates uniformly around an initially bare bead. By allowing filaments to break or debranch above a certain tensile force or branch angle threshold, we found that these networks can crack open, leading to symmetry breaking and bead propulsion (Fig. 7B; Movie S9) as we observed in the experiments (Fig. 1C). To account for changes in CP concentration, we varied the average filament branch length in the simulations.

**Figure 7.**
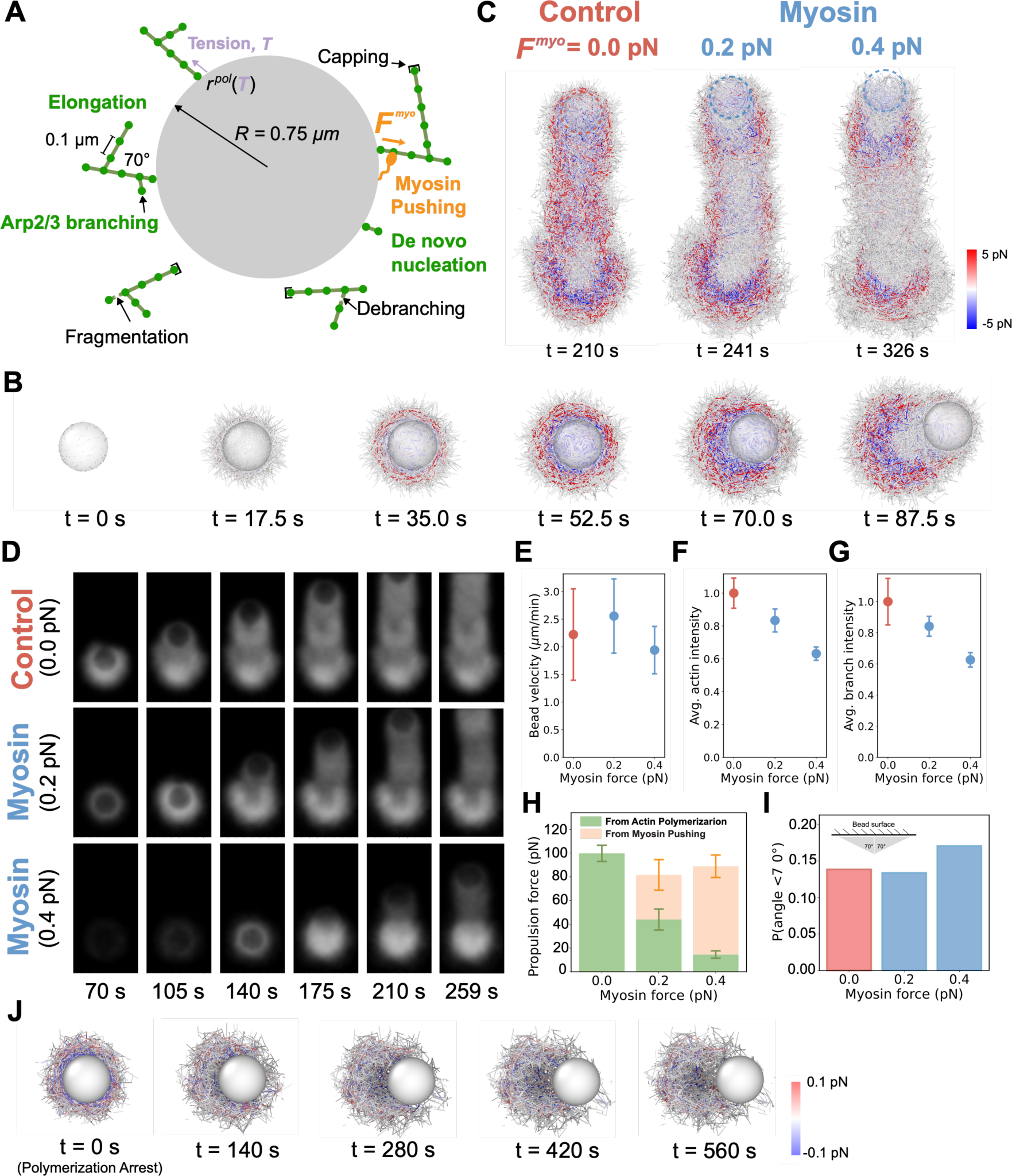
Filament level model of actin comet tail recapitulates experimental results. (**A**) Schematic of model of actin polymerization around the nucleating bead. Model includes filament level nucleation, branching, filament fragmentation, debranching, capping, and force exerted by implicit myosin. (**B**) Timelapse of symmetry breaking event under intermediate capping condition (branch length = 0.5 μm) with no myosin. Color scale indicates filament tension (Red: tensile; Blue: compressive). (**C**) Tension distributions within actin comet tails formed at either control (0.0 pN) or two myosin forces (0.2 pN or 0.4 pN) under intermediate capping condition (branch length = 0.5 μm). Image shows a cut through the center of the comet tail. Color scale indicates filament tension (Red: tensile; Blue: compressive). (**D**) Simulated timelapse of epifluorescence images under different myosin forces (branch length = 0.5 μm). (**E**) Elongation speed (**F**) actin intensity and (**G**) Arp2/3 complex intensity for simulated beads as a function of myosin force (branch length = 0.5 μm). (**H**) Forces acting on beads along the direction of bead propulsion due to actin polymerization (green) and myosin pushing (orange) as a function of the myosin force (branch length = 0.5 μm). (**I**) Orientation of filaments around the beads (within 0.15 μm of the bead surface) as a function of myosin force (branch length = 0.5 μm). (**J**) Simulated symmetry breaking after actin polymerization and branching arrest. Actin was allowed to polymerize around the bead for 42.7 s (0.2 pN myosin force with 0.5 μm filament length) to form a shell of intermediate thickness before halted. The time of arresting was set as t = 0 s.

The effect of the myosin-I power stroke was modeled as constant tangential pushing forces of magnitude *F*^myo^ acting along every actin filament segment close to the bead, with equal and opposite force on the bead (Fig. 7A). With myosin-I pushing force incorporated, simulations recapitulated many of the experimental findings. For the short filament scenario (high CP concentration), simulated myosin-beads showed a significant delay or absence of symmetry breaking due to myosin pushing short filaments away from the bead surface (Fig. S5A, C; Movie S10). For the long filament scenario (low CP concentration), simulations reproduced the accumulation of a dense actin shell around the beads, which significantly inhibited the symmetry breaking of the control-beads (Fig. S5B, C; Movie S11). Myosin-beads by contrast promoted the growth of an asymmetric dense cloud (Fig. S5B, C), similar to the experimental observations in early stages prior to bead ejection (Fig. 2B). We note that simulations do not account for depletion of bulk actin occurring during late stages of bead ejection in experiment (Fig. 2B). We also note a faster accumulation of actin in the simulations compared to the experiments (Fig. S5B), which suggests additional factors limiting network growth not considered in the model. For the intermediate filament length scenario (i.e. intermediate CP concentration), simulations showed a similar elongation speed for the myosin-bead compared to the control-bead, albeit with a less dense actin network (Fig. 7D-F; Movie S12), in agreement with the experimental observations (Fig. 3A-F). Further, simulation predicted that actin filaments in the myosin comet tail experienced less stress (Fig. 7C), possibly due to the sparse actin organization, which reduces stresses arising from piling interconnected actin filaments on top of each other.

Notably, the ratio of actin to Arp2/3 complex density in simulated comet tails did not increase in the presence of myosin (Fig. 7F-G; Fig. S6A), as observed in experiments (Fig. 4D). This fixed ratio is a consequence of our fixed-filament-length assumption to mimic a certain CP concentration. This assumption validates when capping rate reduces equally as actin polymerization rate in response to opposing force, as reported by Li et al (*15, 19*). However, the mismatch between our simulations and the experiments suggests an inequivalent effect on actin polymerization and CP capping due to the pushing force of myosin. We also tested whether the presence of myosin would promote actin debranching, which could also change the actin to Arp2/3 ratio. We found that myosin forces do not enhance debranching in our simulations (Fig. S6). Indeed, less debranching occurs since comet tail stresses are smaller in the presence of myosin (Fig. 7C).

We next dissected the force contribution from different sources (actin polymerization or myosin power stroke) that powered the bead propulsion in our simulations. Strikingly, we found that while in control-beads the symmetry breaking and net comet propulsion were exclusively powered by actin polymerization, in myosin-beads, the myosin pushing forces contributed significantly to bead propulsion after symmetry breaking, and had a predominant effect at high *F*^myo^ (Fig. 7H; Fig. S7). Simulated myosin forces also partly reorganized the network such that a larger fraction of actin filaments was polymerizing facing more perpendicular to the bead surface at larger myosin forces (Fig. 7I). When actin polymerization was arrested before shell breaking, simulations replicated the experimental findings that myosin pushing forces alone can eject the bead out of an actin shell (Fig. 7J; Movie S13), provided the shell is not too dense or too sparse (Fig. S8)

Overall, our simulations with the myosin-I power stroke incorporated as an effective repulsive force replicate most of our experimental observations and support the experimental finding that myosin-I enhances force generation during branched actin assembly to promote shell breaking and enhance the efficiency of bead propulsion.

## Discussion

### Myosin-I force generation and modulation of actin network density

The primary finding of this study is that Myo1d synergizes with Arp2/3 complex to enhance the pushing forces of branched actin networks, and that this force enhancement requires the myosin power stroke. Importantly, we also found that Myo1d alters the actin network composition grown from NPF beads, producing a less dense actin network with decreased incorporation of Arp2/3 complex.

How does myosin-I modulate Arp2/3 complex incorporation? We propose a model where myosin-I motor activity pushes the actin filaments away from the bead surface, reducing the accessibility of actin binding sites for NPF-activated Arp2/3 complex to bind and nucleate new branches (Fig. 8). Alternatively, the mechanochemistry of myosin-I may differently affect the incorporation of actin monomer and CP to filament ends in a way that promotes actin elongation while slowing capping (*19, 42*). Additional experiments that measure the actin elongation rate and capping kinetics near the NPF-coated surface are required to test this hypothesis.

**Figure 8.**
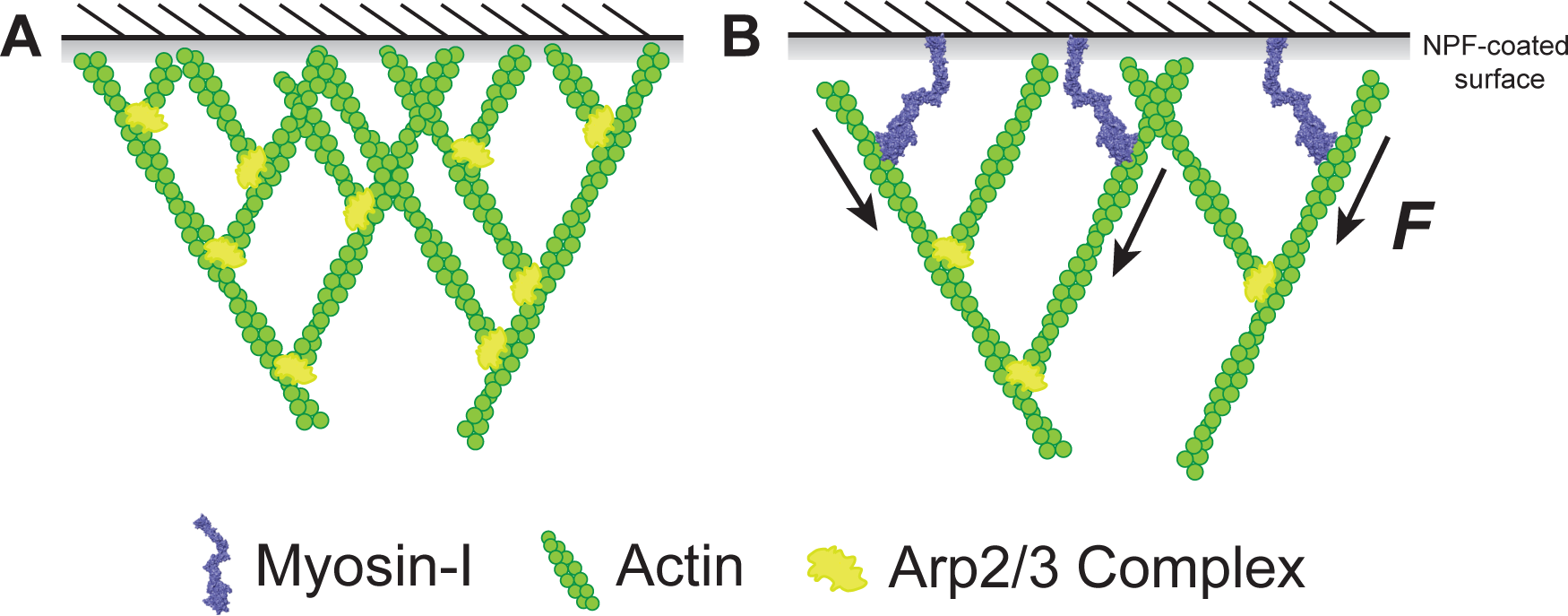
Schematic of how myosin-I modulates actin network structure through its power-strok.

We do not favor a model in which myosin-I sterically inhibits NPFs from binding and activating Arp2/3 complex for three reasons. First, the concentration of SNAP-Arp2/3 complex bound on beads is unchanged in the presence of myosin (Fig. 4E). Second, networks grown in the presence of calcium where the myosin power stroke was uncoupled from actin binding showed similar actin densities between myosin- and control-beads (Fig.5 D, F), which ruled out the possibility that the sparse actin organization was due to the competition between myosin-I and Arp2/3 complex for actin binding sites. Finally, calculations of molecular occupancy on bead surface through protein quantification verified that the steric hindrance effect of myosin-I is likely negligible (See SI for detailed quantifications).

It remains possible that force generation by myosin-I debranches actin filaments, resulting in less dense network with reduced Arp2/3 incorporation. Mechanical forces ranging from 0 pN to 2 pN dissociate actin branches from their mother filaments together with Arp2/3 complex (*43*). A myosin-I paralog (Myo1b) that generates such forces has been shown to dissociate branches via motor activity (*44*). Although our simulations disfavor this hypothesis (Fig. S6), a role for myosin-I induced debranching should be explored further in the cell.

Finally, it will be intriguing to explore these models further by performing experiments and computational modeling to probe network growth under mechanical load in the presence of myosin-I, especially given previous work that demonstrates the substantial loading effects on network architecture and power generation (*15, 19*).

### Myosin-I Cell Biology

Myosin-Is connect the actin cytoskeleton to cellular membranes where they contribute to plasma membrane dynamics, organelle deformation, and shaping of actin network architecture. Although the molecular details of myosin-I function have been difficult to determine, recent studies suggest some paralogs are powerful motors that have active roles in shaping membranes. For example, myosin-Is in budding yeast have mechanochemistry suitable for generating power working with polymerizing actin to drive membrane invagination during endocytosis (*23, 24, 45*). Our current study confirms the ability of myosin-I to exert substantial pushing forces capable of fracturing actin networks. This power-generating capacity likely translates across species, with vertebrate Myo1c regulating actin architecture in diverse cellular regions (*46–48*) and Myo1e playing key roles in processes like endocytosis (*49, 50*), phagocytosis (*26*), and cancer invadosome formation (*51, 52*). Taken together, these findings point to some myosin-Is as dynamic players, actively shaping cellular structures and processes through their unique ability to both link and manipulate membranes and the actin cytoskeleton.

Not all paralogs are expected to modulated the actin networks as Myo1d. Notably, some myosin paralogs have substantially slower kinetics which are better suited for a motor that functions as dynamic tether, providing force-dependent linkages between actin and membranes (*22–24, 45, 48, 53, 54*). Our experiments performed at low ATP concentrations (Fig. 5 A-C) mimic the behavior of myosins with slow motility rates, and show clearly that slow kinetics can inhibit network fracturing and comet tail growth. Further studies to investigate how the intrinsic kinetic properties of different myosin-I paralogs influence actin polymerization and membrane dynamics will further reveal the diverse role of myosin-I function (*23, 24, 26, 45, 48, 54*).

### Summary

Overall, our study provides insights into how myosin-I molecules coordinate with Arp2/3 complex, regulating the dynamics and architecture of the branched actin network and promoting the actin-based motile force generation (Fig. 8). This work sheds light on synergy between myosin motor activity and actin polymerization, underscoring their collective role in driving morphological changes at the cellular membrane interfaces. Future work to determine how myosin-I affects actin network architecture and polymerization forces will reveal the molecular roles of this important myosin family.

## Materials and Methods

### Protein Purification

Actin was purified from rabbit skeletal muscle acetone powder as previously described (*55*). Monomeric G-actin was purified by gel filtration on Sephacryl S-300 in G-buffer (2 mM Tris–HCl [pH 8.0], 0.2 mM ATP, 0.1 mM CaCl2, 1 mM NaN3 and 0.5 mM DTT) and used within 2-3 weeks. Actin was labeled with NHS-Rhodamine at random surface lysine residues (*56*). Full-length *Drosophila* Myo1d, with C-terminal FLAG-Avi tags, was expressed, purified as previously described (*57*), and subsequently biotinylated at the C-termini Avi tag sequence via BirA biotin-protein ligase (Avidity) (*31*). GST-tagged WCA domain of human WASP protein (GST-WCA) was purchased from Cytoskeleton (Cat. #VCG03-A) and used without further purification. Arp2/3 complex was isolated from bovine brain as previously described (*58*). SNAP-tagged Arp2/3 complex was constructed and purified as described (*59*). SNAP-tagged Arp2/3 complex was labeled with SNAP-Surface 488 (Biolab, Cat. # S9124S) using commercially provided protocol. Human CapZ was expressed and purified as previously described (*60*). Halo-ABD was constructed and purified as described in (*61*).

### Bead Preparation

Carboxylate polystyrene beads (Polybead, 2.0 μm diameter, CAT. #18327-10) were purchased from Polysciences. Beads were coated with NPF and neutravidin following a previous protocol (*28*) with modifications. Briefly, 5-10 μL bead slurry was washed with X buffer (10mM HEPES [pH7.5], 100 mM KCl, 1 mM MgCl2, 100 μM CaCl2, 1 mM ATP), and then incubated with 50-100 μl 2.3 μM (0.1mg/ml) GST-WCA and various concentrations of Neutravidin (ThermoFisher, Cat. #31000) (8.3 μM (0.5mg/ml) for #myosin/NPF=0.28:1, 16.7 μM (1mg/ml) for #myosin/NPF=0.35:1, 33.3 μM (2mg/ml) for #myosin/NPF=0.43:1, or 83.3 μM (5mg/ml) for #myosin/NPF=0.80:1) on a slow rotator at 4 °C for 2hr. Beads were then pelleted by spinning at 16000 g for 2 min at 4 °C, to remove the unreacted reagents, and then resuspended in 200-400 μL 10mg/mL BSA, incubating on ice for 30 min to block the free space left on bead surface. Finally, beads were washed twice and stored in 50-100 μL 1mg/mL BSA in X buffer for up to 3 days.

The NPF and neutravidin-coated beads were next split in half, with one half coupled with biotinylated *Drosophila* Myo1d (myosin-bead) or biotinylated Halo-ABD, and the other half coated with Biotin-CF640 (Biotium, Cat. #80032) fluorescence dye (control-bead). The NPF and neutravidin-coated beads were first washed with M buffer (20 mM HEPES [pH7.5], 100 mM KCl, 1 mM MgCl2, 1 mM EGTA, 2 mM ATP) to remove the free calcium in the storage X buffer, and were then incubated with 1 μM biotinylated *Drosophila* Myo1d, 1 μM Halo-ABD, or 1 μM Biotin-CF640 fluorescence dye for 30 min on ice. Beads were then pelleted under 16000xg for 2 min at 4 °C, to remove the free unbound reagents, and washed with 1mg/mL BSA in M buffer and used immediately for motility assay on the same day after preparation.

### Bead motility assay/comet tail assay

Unless specified otherwise, a typical motility mixture contained 4 μM actin (5% Rhodamine labeled), 200 nM Arp2/3 complex or SNAP-Arp2/3 complex, 6.5 - 200 nM CP, 2 μM Calmodulin, and 3 μL bead slurry (1.5 μL of myosin-beads or HaloABD-beads, and 1.5 μL of control-beads), mixed in 20 mM HEPES [pH7.5], 100 mM KCl, 1 mM MgCl2, 1 mM EGTA, 1 mM MgATP, 40 mM DTT, 10 mg/mL BSA and 0.2% methylcellulose, contributing to a final volume of 50 μL. The activity of SNAP-Arp2/3 complex is slightly lower than the unlabeled complex, so we changed the CP concentration (40 nM) to achieve similar tail lengths, actin densities and growth efficiencies as observed for the native complex (Fig. 4). For experiments with calcium present, G-actin was pre-incubated with 200 µM EGTA and 50 µM MgCl2 for 5 min to be exchanged to Mg-G-actin before using. Calcium experiments included 1.1 mM CaCl2 in place of Calmodulin to disrupt the myosin power stroke. Beads were first mixed with all other reagents, excluding actin, with a pipette, to ensure an even distribution. The reaction was then initiated by adding actin into the system, mixed thoroughly, and denoted as time 0. Slides and coverslips were wiped with 70% ethanol and ddH2O, followed by a plasma cleansing for 10 min. Upon mixing, a volume of 2.1 μL motility mixture was carefully applied between a glass slide and a coverslip (22 mm × 22 mm) forming a so called ‘squeeze chamber’ with a height of 4.3 μm. The squeeze chamber was then sealed with vacuum grease and imaged immediately under the microscope. Time-lapse movies were acquired of microscope fields that included both myosin- and control-beads. In most cases, the actin comet tails emerging from myosin- or control-beads grew with a constant speed during the first ∼10 minutes following symmetry breaking. As the reagents in the polymerization mixture depleted, tail elongation slowed and eventually stopped ∼30 min after mixing.

### Fluorescence imaging and Data analysis

Fluorescence microscopy imaging was performed via Leica DMIRB epifluorescence microscope (100x, oil-immersive objective of numerical aperture 1.4) and Metamorph (Molecular Devices) imaging software. Movies were recorded at 25 °C and acquired every 10 s for 30-60 min. Exposure time was 200 ms for most experiments, and 1s to image the SNAP-Surface 488 labeled Arp2/3 complex. Images were analyzed and quantified using Fiji software. Actin comet tail length were measured manually at each frame using the segmented line draw tool (from the end of the tail to the center of the bead) and converted to the microns. Tail growth rate and fluorescent assembly rate were calculated by fitting the first 7 to 10 data points in the time courses to get the initial slope of the growth and assembly. Growth efficiency was determined by dividing the tail growth rate by the fluorescent assembly rate. Image brightness and contrast were carefully adjusted using Fiji and Adobe Illustrator. Unless otherwise specified, the lookup table for each pair of control- and myosin-bead (or HaloABD-bead) were kept the same.

### Statistical analysis

The statistical significance was calculated using paired or unpaired two-tail Student’s t test in Prism v9.0. Further details are described in figure legends.

### Modeling methods

We simulated actin comet tail formation at the level of individual filaments, each represented as a series of segments of length *l*_0_, following earlier work (*62*). The pointed ends of actin filament branches are assumed to be connected to a point element of a mother filament, at the location of Arp2/3 complex. Such a filament representation allows us to model the effect of myosin as a tangential force acting along filament segments close to the nucleating bead. We thus generalize earlier filament models that did not explicitly account for filament bending mechanics (*35, 63–65*), earlier mechanical models that did not monitor the whole network of actin filaments (*36, 66–69*), or modeled the full process of symmetry breaking and propulsion (*70*). We do not explicitly consider diffusiophoretic contributions to bead motion (*71*). (Also see SI for further details).

#### Forces on actin filaments

The position ***r_i_*** of point element *i* of actin filaments/filament branches evolves according to (*62*)

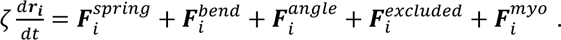

Here *ζ* is an effective filament segment drag coefficient that allows the actin network to evolve through approximate quasi-static mechanical equilibrium, while also providing numerical stability. The spring force is 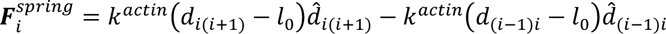 where (*i*-1), (*i*+1) are neighboring point elements (if they exist) before and after *i, d_ij_* is the separation distance between *i* and *j,* and 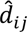 is the unit vector from *i* to *j*. The equilibrium length is *l*_0_, except for (i) uncapped barbed ends that elongate on average according to 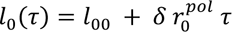 after their initiation at *τ* = 0, in discrete steps of half-monomer size *δ* = 2.7 nm (see below) and (ii) a short branch segment joining the mother filament point element to the daughter pointed end, which has length of *l_branch_*.

The bending force is 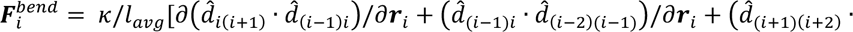,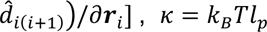 is the flexural rigidity, *l_p_* is the persistence length of the actin filament, and *l_avg_* is the average length of the two filament segments composing the angle. For the straight angle that exists among the first three beads in the connection between mother and daughter filaments *l_avg_* = *l*_0_ for numerical stability.

An angular potential keeps Arp2/3 complex branches at 70°. The angular force is 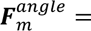,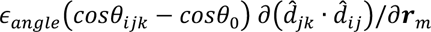 where *ϵ_angle_* is a spring constant, *i, j, k* are the indices of the point elements that make up the angle, and *m* is one of these indices.

The excluded volume force is due to repulsion between two actin filaments and between actin filaments and the nucleating bead. The former prevents crossing of filaments and is exerted along the direction of vector ***d_αβ_*** that connects the two closest approach points on the filament segments α and β (*72*). It is modeled as a stiff spring force with a max range of *d^excluded^*, with 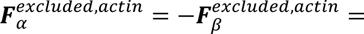,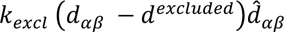, where *d_αβ_* is the minimum distance between neighboring filament segments. This force is distributed to the endpoints of filament segments α and β (including element *i*) according to a level arm rule. Though the typical diameter of actin filaments is around 7 nm, we set *d^excluded^* to be 20 nm in order to mimic the effect of thermally fluctuating filaments taking up a larger volume (which is not included in our simulations). Additionally, since the excluded volume is modeled as a stiff spring rather than a hard-core potential, some level of overlap between segments is possible. At a separation of order the filament diameter, 7 nm, the excluded volume force is 22 pN. Filament segments experience a radially-oriented excluded volume force with the nucleating bead if they are closer to the nucleator than *R*; this force can be written as 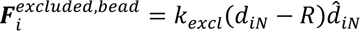 where *d_iN_* is the closest approach distance between the filament segment and the nucleator bead.

Myosin forces are given by 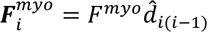, where *i*-1 is the neighboring actin filament point element along the pointed end direction. It acts on all filament elements *i* closer than *d^myo^* to the nucleator bead surface. The myosin force acts tangentially along the segment towards the pointed end. We do not model the individual binding, lever arm motion, and unbinding of myosin; instead the magnitude of *F^myo^* approximates a time and ensemble average over many binding and unbinding cycles.

#### Nucleating bead motion

The nucleating bead evolves through time according to:

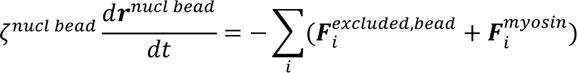

Where the sum is over all actin segments that contact the bead through excluded volume or myosin interactions, according to the interaction distances defined above. We assume the relative motion between actin network and nucleating bead is dominated by frictional forces between them. We used a value of *ζ^nucl bead^* that was large enough to prevent the rapid ejection of bare beads out of a shell at the onset of symmetry breaking, a phenomenon that is not seen in our experiments. We thus monitor the relative motion of the bead and actin network in the limit of quasi-static mechanical equilibrium of the actin network, including implicit transient attachment and detachment of actin filaments to the bead. This approximation does not account for the varying concentration of actin near the bead or the absolute motion of actin comet and bead in the lab frame, which in reality would be influenced by small forces between the bead or actin and the glass slide.

#### Actin barbed end polymerization rate

Polymerization of the barbed end of uncapped filaments occurs with the last segment of the actin filament lengthening in increments of half-monomer size *δ* at a rate given by the polymerization rate. If the end segment reaches a length of *l*_0_, a new segment is added with initial length *l*_00_. Polymerization of the barbed end is attenuated by compressive forces on the spring bond connecting the barbed end point element to the rest of the filament. The polymerization rate is 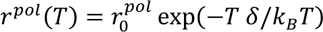, where the free filament elongation rate is 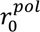 and 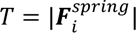 when 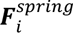 is the compressive tension on the barbed end point element *i* (otherwise 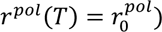 (*73, 74*). Because each filament segment starts with an initial length of *l*_00_ (for numerical stability), the rate *r^pol^*(0) is effectively multiplied by a factor of 1.12.

#### Barbed end capping

Filaments polymerize until they reach a final length specified for a given simulation, *L_fil_* (an integer multiple of *l*_0_), which we vary to simulate the effect of varying CP concentration. Here we didn’t study the effects of a varying filament length distribution. The assumption that *L_fil_* is independent of the force is based on prior experiments (*19*) and the fact that the filament length added by capping protein is close to *δ*. According to this evidence, the polymerization to capping rate ratio, as well as the average filament length, remains unchanged by force for a given CP concentration.

#### Branching

Branching occurs from actin filament point elements that are within *l*_0_ of the nucleating bead, at rate 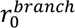, independent of the CP concentration. The rate was chosen to approximate the timescale of symmetry breaking and comet speed at intermediate CP. The segment length connecting the mother filament point element to the daughter filament pointed end point element is *l^branch^*. Their orientation is chosen according to a uniform angle distribution in the cone around the mother filament opening towards the barbed end. To maintain a discretization of the network at segment length *l*_0_, branches cannot branch from the barbed end point element of the filament. Branches nucleating from the same mother filament have no torsional restriction (i.e. daughter filaments can rotate about the axis of the mother filament without restriction), but implicit torsional restrictions are imposed due to the dense surrounding network limiting this motion.

#### De novo nucleation

Nucleation of new filaments of length *l*_00_ occurs at a rate of 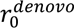. These small filaments are introduced with a uniform spherical orientation with either their pointed or barbed end touching the nucleating bead surface. This slow rate of filament introduction was tuned to allow for startup of the network on timescales comparable to experiment, and to allow for the buildup of a thin actin cloud around the leading bead edge during actin comet propulsion, as observed in experiments. At the beginning of the simulation, 200 of these de novo filaments are added to provide enough initial filaments for symmetric growth of the shell.

#### Filament fragmentation and debranching

For simplicity, filament fragmentation was assumed to occur above a certain tensile force threshold *F^frag^* (instead of implementing a rate of severing on as a function of tension).

For filament segments the threshold was *F^frag^* = 50 pN. This is lower than the experimental fragmentation force of phalloidin-labeled actin filaments, which is several hundred pNs (*75*), but we found that the shell would have difficulty breaking unless we had this lower force. Once a filament segment is fragmented, the filament with the newly created barbed end is left uncapped and can grow to the final length specified for that simulation. Debranching occurs for branches that have a branch bond with tension greater than *F^debranch^* = 30 pN or if the angle deviates by more than *Δθ^debranch^* = 25° off of the 70° equilibrium angle. The value of *Δθ^debranch^* was chosen to allow easy debranching when the branch is bent (similar to experiments where branches were bent and pulled by fluid forces of order pN, (*43*)). The value of *F^debranch^* had to be sufficiently high such that the network does not easily fall apart.

## Acknowledgments

We thank Daniel Safer, Faviolla A. Baez-Cruz and Richard Wike for their kind help with protein purifications and technical assistance on the project. We thank everyone in Ostap Lab and Dominguez Lab for their valuable inputs throughout the project duration.

## Author contributions

Conceptualization: M.X., D.M.R., L.W.P., R.D., D.V., E.M.O.

Methodology: M.X., D.M.R., L.W.P.

Formal analysis: M.X., D.M.R.

Investigation: M.X., D.M.R., L.W.P.

Resources: G.R., M.B.

Visualization: M.X., D.M.R. L.W.P., D.V., E.M.O.

Supervision: L.W.P., R.D., D.V., E.M.O.

Writing—original draft: M.X., D.M.R., D.V., E.M.O.

Writing—review & editing: M.X., D.M.R., L.W.P., M.B., R.D., D.V., E.M.O.

## Data and materials availability

All data needed to evaluate the conclusions in the paper are present in the paper and/or the Supplementary Materials. Additional data related to this paper may be requested from the authors.

## Supplementary Text

### Myosin steric hindrance effect

We quantified the average spatial occupancy of individual components (NPF or myosin-I) bound to bead surface for the first three myosin/NPF ratio groups (0.28, 0.35, 0.43), all of which maintain the same NPF densities (∼ 6000 /um^2^). We found that individual molecular components occupied similar areas in all three groups: 11.4×1.4 nm^2^/molecule for 7680 molecules in 0.28:1 (myosin/NPF) group, 11.1×11.1 nm^2^/molecule for 8100 molecules in 0.25:1 (myosin/NPF) group and 10.7×10.7 nm^2^/molecule for 8580 molecules in 0.43:1 (myosin/NPF) group. Therefore, the steric hindrance effect in each of the three groups should be nearly identical. In the 0.28:1 (myosin/NPF) group, both control- and myosin-beads displayed similar actin comet patterns, which implies that the steric hindrance of myosin-I does not play a significant role in modifying the Arp2/3-medicated branching under the current conditions.

### Bead gliding assay on actin filament tracks

F-actin was made by polymerizing 25 μM G-actin in MB buffer (10 mM MOPS [pH7.0], 25 mM KCl, 1 mM EGTA, 1 mM MgCl2, 1 mM DTT) with 2 mM ATP at room temperature for 1 hr, and stabilized with AlexaFluor 488 Phalloidin (ThermoFisher Cat. #A12379). Myosin-beads were prepared following the procedures outlined in Methods in the main text and labeled with TRITC during the blocking process using 10% TRITC-labeled BSA (Sigma, Cat. #A2289). Flow chambers were made by sandwiching the plasma cleaned slides and coverslips (22 mm × 22 mm) with two pieces of double-stick tape, forming a 5mm-wide channel that can hold a volume of up to 10 μL. To immobilize single-actin-filament tracks on the coverslip, we coated the flow chamber with 0.5 mg/mL NEM-myosin, incubated for 5 min, followed by two washes with 2 mg/mL casein (1 min for each) to remove any unbound NEM-myosin and block the remaining free spaces on the surface. Next, 5-10 nM AlexaFluor 488 Phalloidin stabilized F-actin was loaded into the flow chamber to avoid any overlapping between the actin filaments, incubated for 2 min, followed by a wash with 20uL MB buffer. Finally, 10 uL of the 50 uL bead motility mixture containing 3uL TRITC-labeled myosin-bead slurry (#myosin/NPF=0.4:1), 2 μM Calmodulin, 0.2 mg/mL glucose oxidase, 0.04 mg/mL catalase and 5 mg/mL glucose in MB buffer supplemented with 5 mM Mg-ATP and 40 mM DTT, was loaded into the flow chamber, sealed with vacuum grease and imaged immediately under the microscope.

### Myosin inhibition experiments

Rigor myosin was prepared by excluding ATP from the motility mixture. Gel-filtered actin monomers were incubated with 2 mM ATP for 15 minutes on ice, followed by passage through a G25 Sephadex column to remove residual free nucleotides. The resulting ATP-treated actin exhibited similar polymerization capability in ATP-free mixtures as to those containing 1 mM ATP. Alternatively, myosin was inhibited by introducing 10 mM Mg-ADP to prolong the lifetime of the myosin ADP-bound state.

### Actin gliding assay

Coverslips were plasma-cleaned and coated with 0.5% Nitrocellulose and used to make a flow chamber. Flow chamber was first incubated with 0.1 mg/mL neutravidin for 2 min followed by blocking with 2 mg/mL casein, twice, 1 min each. Then, 250 nM biotinylated *Drosophila* Myo1d was loaded into the chamber, incubated for 2 min, and washed with 20uL MB buffer (10 mM MOPS [pH7.0], 25 mM KCl, 1 mM EGTA, 1 mM MgCl2, 1 mM DTT). Next, 20 nM F-actin stabilized with AlexaFluor 488 Phalloidin was loaded into the chamber, incubated for 2 min, washed with 20uL MB buffer. Finally, the imaging mixture containing 8.4 μM Calmodulin, 1 mM Mg-ATP, 40 mM DTT in 20 mM HEPES [pH7.5], 100 mM KCl, 1 mM MgCl2, 1 mM EGTA and 0.2% methylcellulose was loaded into the chamber and observed immediately. For experiments with calcium, 8.4 μM Calmodulin was replaced by 1.1 mM CaCl2 to disrupt the myosin power-stroke.

### Actin arresting

Actin assembly was quenched by combining equal volume of motility mixture and arresting solution, which contains 20 μM (5 molar excess) of both Latrunculin B and CK-666 (together with 20 μM phalloidin for ‘+ Phalloidin’ experiments) in 20 mM HEPES [pH 7.5], 100 mM KCl, 100 μg/ml BSA, 0.5 mM TCEP. Quenched motility mixture was either applied on a squeezer chamber or loaded into a NEM-myosin coated flow chamber (to make sure the observed shell-breaking was not due to the squeezing effect between the slides and the coverslips) and imaged under the microscope immediately after sealing with vacuum grease.

### Computational modeling

#### Reduction of de novo and branch nucleation rate due to crowding

If upon introduction to the system a de novo filament or new branch would strongly overlap with other filaments or the nucleating bead, their insertion is rejected. Specifically, their introduction is rejected if the closest distance between a newly added segment and an existing segment is less than /4, or if the distance between the newly added branch and the nucleating bead is less than (*d^excluded^*/2 + *R*)/4. No explicit reattempts at introduction of a rejected filament are performed, and the rate of branching, 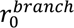, and the rate of de novo introduction, 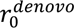, are left unchanged by a rejection event.

#### Simulation timescale

We wrote C++ code with force parallelization using OpenMP. For the case of Fig. 7B (0.5 μm length filaments with no myosin force) it takes 360 hours on 10 CPU cores to run the simulation to 260 s. The amount of time to run a simulated second increases as the simulation proceeds since no filament segments (beyond those that are severed) are removed from the system. Simulations with myosin or with shorter filaments take a shorter amount of time to run the same simulated time since they have overall less filament segments at the same point in time.

#### Simulated fluorescence images

To generate simulated fluorescence images of actin comet tail formation we first rotate the system so the net displacement of the nucleating bead occurs along the x-direction. For a given frame, each actin or Arp2/3 complex spring bond centroid is placed in a 2d grid with gridlines along the x and y axes and bin size 0.05 μm (z-component is not considered). Actin filament segments of length *l*_0_ contribute 37 actin monomers to the bin they are assigned, while shorter filament segments contribute 37 *l*/ *l*_0_monomers, where *l* is the length of the segment. Arp2/3 complex bonds contribute a single Arp2/3 molecule to the bin they are placed in. A grayscale image is constructed from the grid where each voxel in the image corresponds to a single bin and the intensity corresponds to the sum of the count of monomers in each bin. Finally, a Gaussian filter with standard deviation 0.1 μm is applied to the image to smooth the image to a comparable level as in experiment.

#### Measurement of Forces

Forces in Figure 7H and Figure S7 are calculated by first rotating the system so the net displacement of the nucleating bead occurs along the x-direction. Excluded volume forces acting between actin segments and the nucleating bead are spatially divided into three regions based on the closest approach point of the actin segment to the nucleating bead. Forces where the closest approach point is greater than R/4 behind the nucleating bead center in the x-direction are considered back forces. Forces where the closest approach point is greater than R/4 in front of the nucleating bead center in the x-direction are considered front forces. All other forces are labeled as side forces. Excluded volume forces are further subdivided into whether they are caused by filament segments at their maximum length *l*_0_ (actin excluded volume) or filament segments with length less than *l*_0_ (actin polymerization).

#### Measurement of filament orientation with respect to nucleating bead

Filament angle distributions with respect to the nucleating bead, for filaments close to the nucleating bead in Figure 7I are calculated by first rotating the system so the net displacement of the nucleating bead occurs along the x-direction. Only filaments segments with length less than *l*_0_ that are also in the back region (as defined in *Measurement of Forces*) and are also closer to the nucleating bead surface than 0.15 μm are considered. For these actively growing filaments we measure the angle between the filament displacement vector and the vector joining the midpoint of the filament segment to the nucleating bead center. This angle is zero if the filament is perpendicular to the nucleating bead surface and 90° if the filament is lying flat on the bead surface. We generate a probability distribution of these angles over approximately 70 s and when the nucleating bead is moving at a relatively constant velocity. The probability of finding the angle to be less than 70° from these distributions is reported in Figure 7I.

**Fig. S1.**
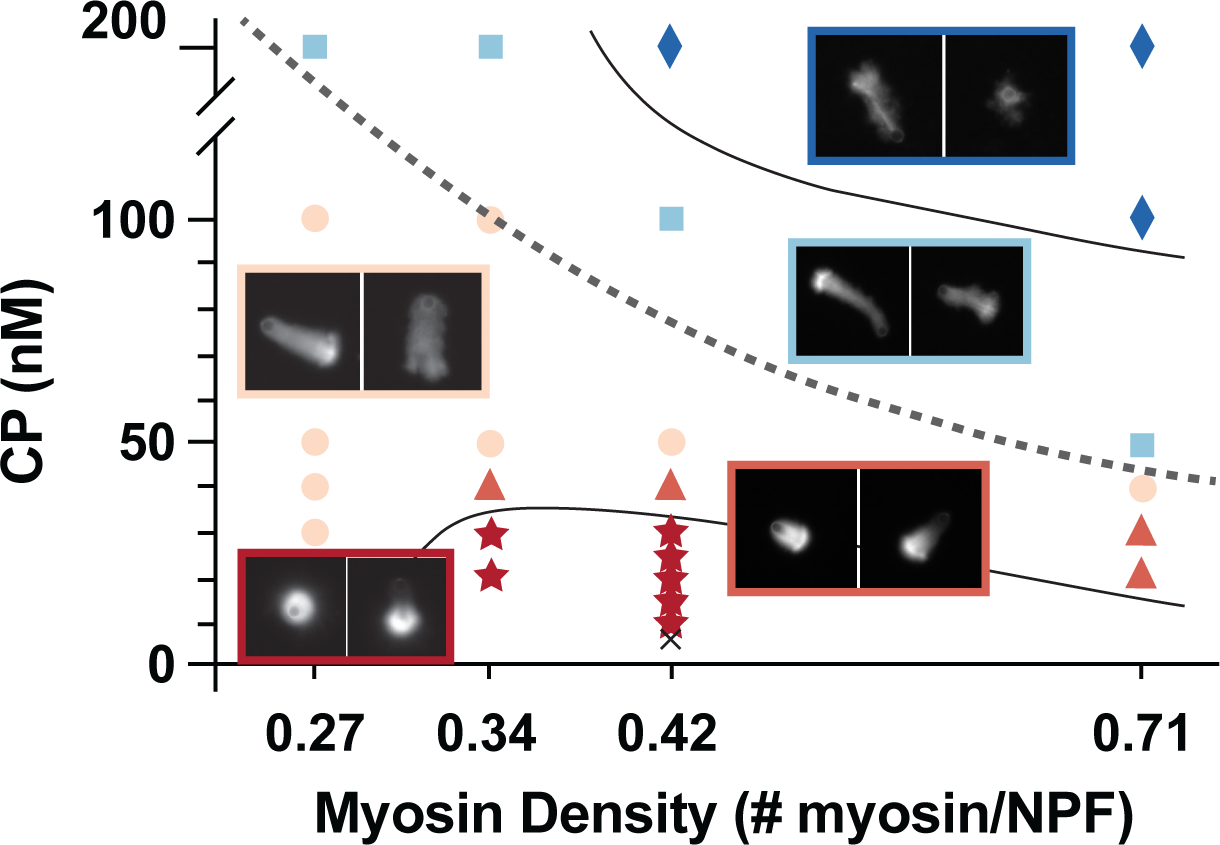
Phase diagram of actin comet tails assembled under different CP concentrations and myosin densities. Experimental points demonstrate control- and myosin-bead pairs assembled under each specified conditions. Star – myosin-beads promote symmetry breaking; Triangle-myosin-beads have longer tails than control; Circle – myosin-beads exhibit similar tail length to control; Square – myosin-beads have shorter tails than control; Diamond – myosin-beads shows no comet tail. For extremely low CP concentration (e.g. 6.5 nM, Crossline), both myosin-coated and control-beads displayed aster-like structure. No significant difference was observed between the two types of beads. Conditions: 4 µM G-actin, 200 nM Arp2/3 complex, 0-200 nM of CP as indicated in graph.

**Fig. S2.**
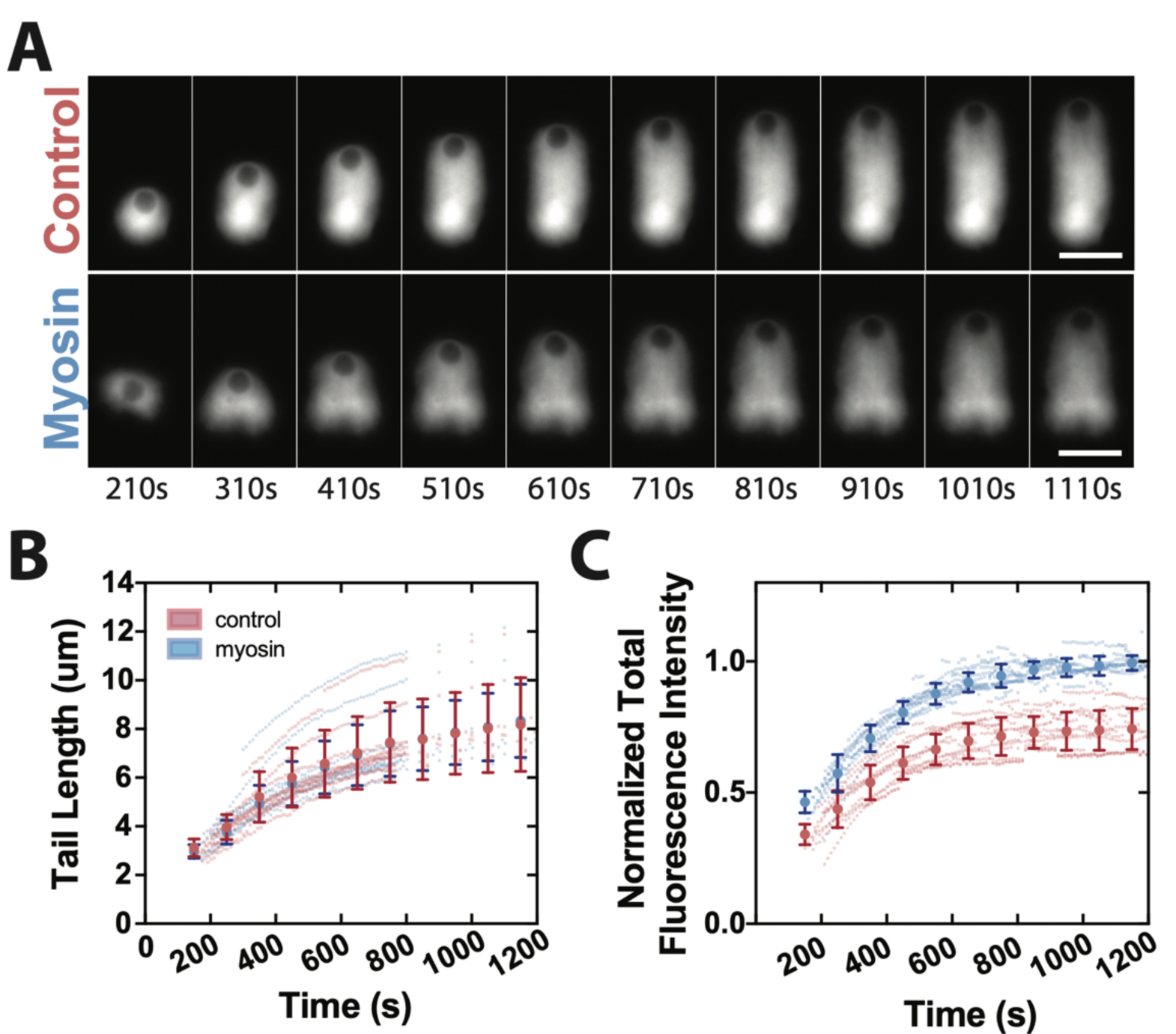
Myosin-I decreases the actin density in comet tails (supplement for Fig. 3 in the main text). (A) Time-series of (top) control- and (bottom) 0.43:1 myosin-beads growing comet tails in the presence of 50 nM CP. The actin network growing from the myosin-bead is less dense but has a similar tail length as the control. (**B**) Actin comet tail length as a function of time (**C**) Comet tail fluorescence as a function of time. The fluorescent intensity of control- and myosin-bead experimental pairs are normalized to the average fluorescence level of the control-beads from 1100-1300 s. Large points show the averaged value at binned time interval (every 100 sec). Traces are from individual beads with each myosin-bead acquired with a control-bead in the same field of view (N=5, n=11). Error bars are SD. Conditions: 4 µM actin (5% Rhodamine labeled), 200 nM Arp2/3 complex and 50 nM CP. Scale bar 5 µm.

**Fig. S3.**
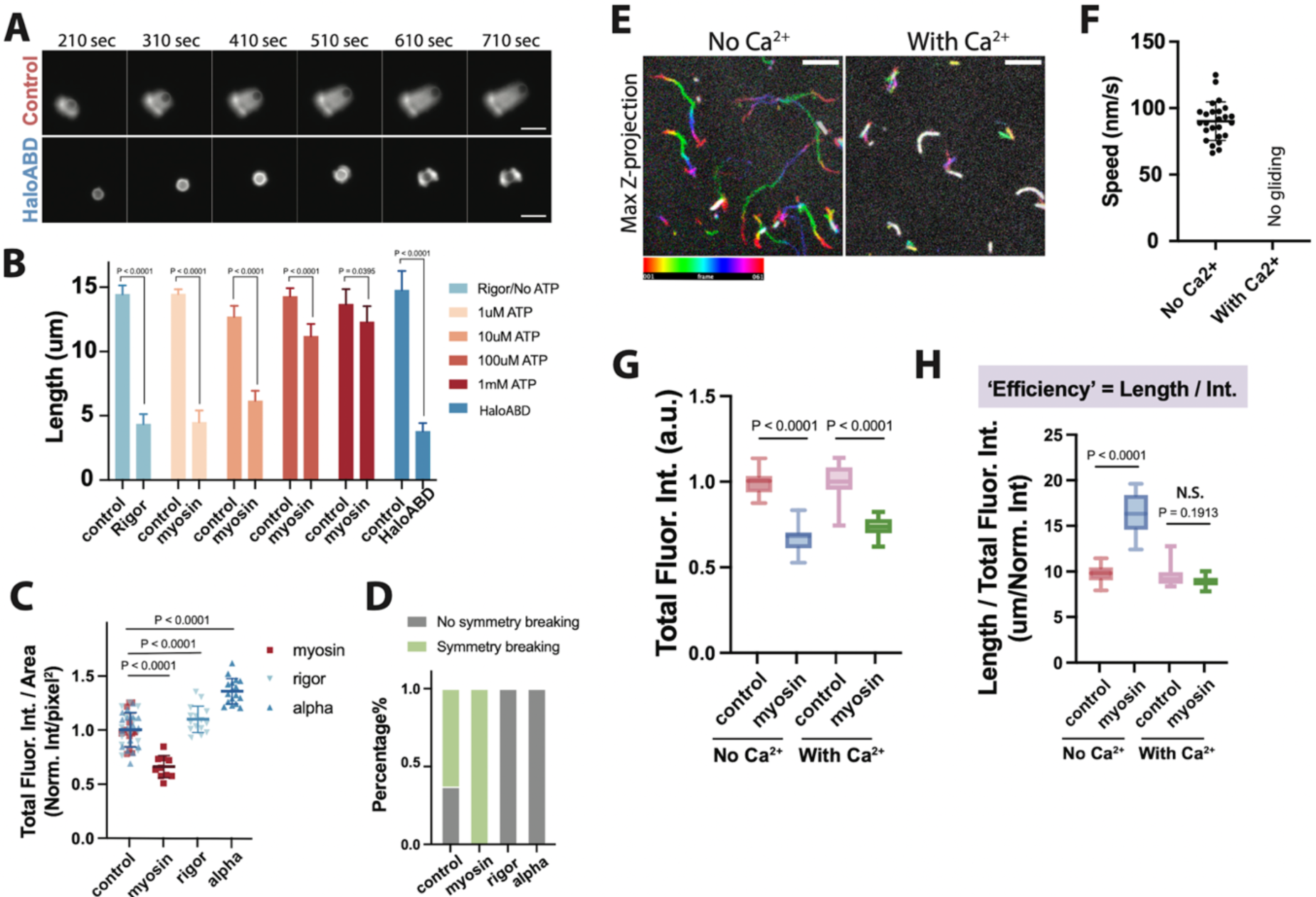
The myosin power-stroke is required for altering network architectures (supplement for Fig. 5 in the main text). (**A**) Time-series of actin assembly around (top) control- and (bottom) HaloABD-bead at 50 nM CP. HaloABD of **α**-actinin heavily delayed the growth of actin comet tail as rigor myosin. Scale bar 5 µm. (**B**) Quantification of comet tail length generated under different ATP concentrations (0-1 mM) for control-, myosin- and HaloABD-beads (N=2, n>15). Error bars are SD. p values were calculated using unpaired two-tail t test. (**C**) Network density quantified by total fluorescence intensity per area for control-, myosin-, rigor-myosin and HaloABD-beads. Normalized to the mean value of control-beads (N=2, n>11). Dot plots show mean ± SD. p values were calculated using unpaired two-tail t test. (**D**) Percentages of control-, myosin-, rigor-myosin and HaloABD-beads that performed symmetry breaking or no symmetry breaking at 15nM CP (N=3, n>23). (**E**) Time collapsed images of actin filaments gliding (with no Ca^2+^) or swirling (with 100 µM free Ca^2+^) on a Myo1d-coated surface. Rainbow bar represents timescale over 61 frames with 5 s frame interval. Scale bar 5 µm. (**F**) Quantification of actin gliding speed with and without 100 µM free Ca^2+^. (**G**) Total actin fluorescence intensity and (**H**) Growth efficiency for control- and myosin-beads with and without the presence of 100 µM calcium (N=1, n=15). Efficiency is defined as comet tail length per unit actin fluorescence intensity. Box plots (**G, H**) show median (center line), interquartile range (box) and min-max values (whiskers). p values were calculated using two-tail paired t test.

**Fig. S4.**
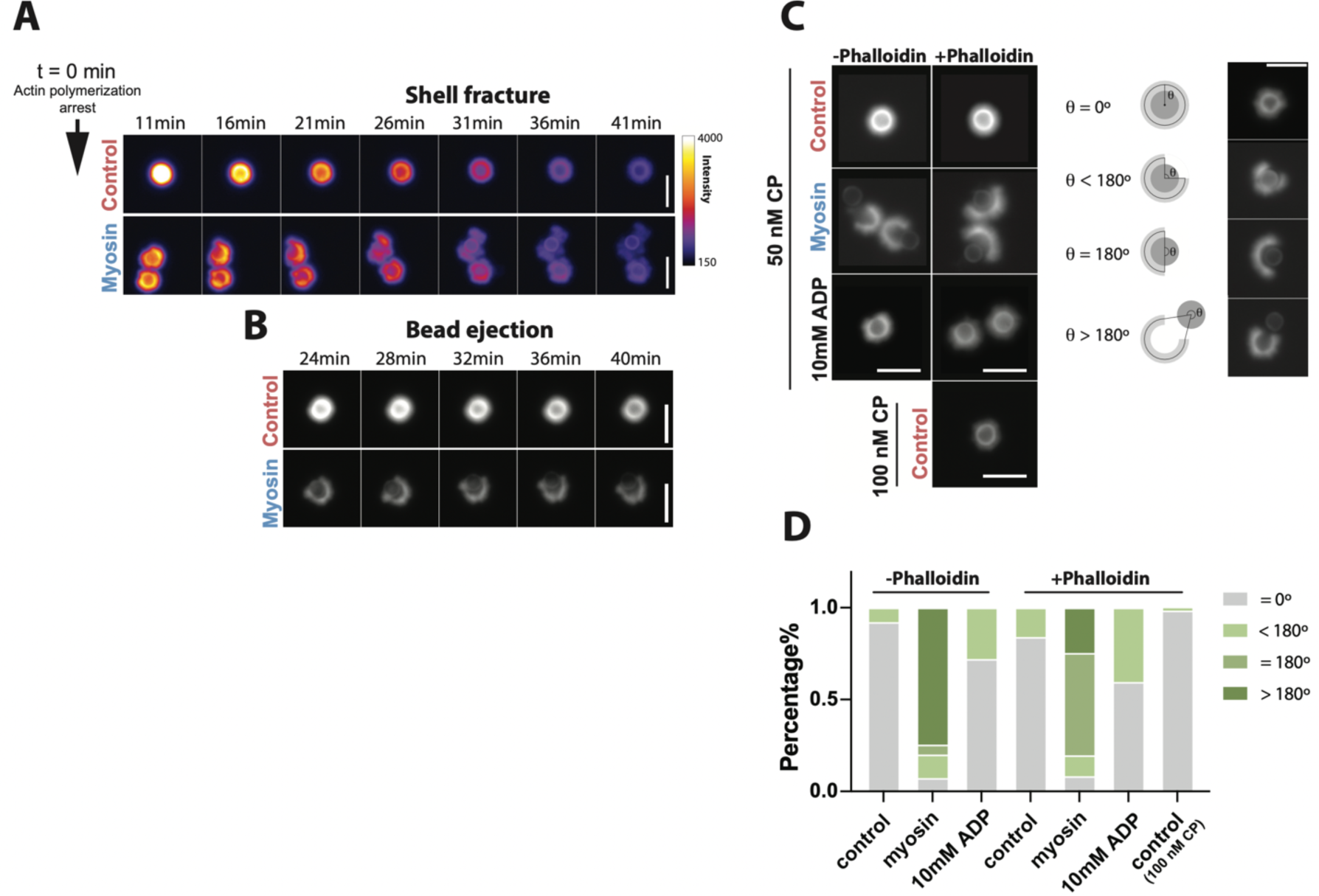
Myosin power-stroke alone can break sparse actin shell (w/o Phalloidin). (A-B).Time-lapse series of (top) control- and (bottom) myosin-bead that performed (**A**) shell fracture or (**B**) bead ejection. Control- and myosin-beads were mixed with 4 μM actin (5% Rhodamine labeled), 200 nM Arp2/3, and 50 nM CP, incubated for 100s, and then actin assembly was arrested by adding 20μM (5 molar excess) of Latrunculin B and CK-666. (**C**) Status of shell breaking approximately 50min after arrest. The extents of shell breaking was classified by shell-breaking angle θ. (**D**) Percentage of different populations with different extents of shell breaking. Without Phalloidin: control (n=141), myosin (n=55), myosin with 10mM ADP (n=125); With Phalloidin: control (n=182), myosin (n=158), myosin with 10mM ADP (n=89). Conditions: 4 μM actin (5% Rhodamine labeled), 200 nM Arp2/3 complex, 50 nM or 100 nM CP was incubated for 100 s and arrested by adding 5 molar excess of Latrunculin B, CK-666 and phalloidin (for ‘+ Phalloidin’ experiments) or 10mM ATP (for ‘myosin with 10 mM ADP’ experiments). Scale bar 5μm.

**Figure S5.**
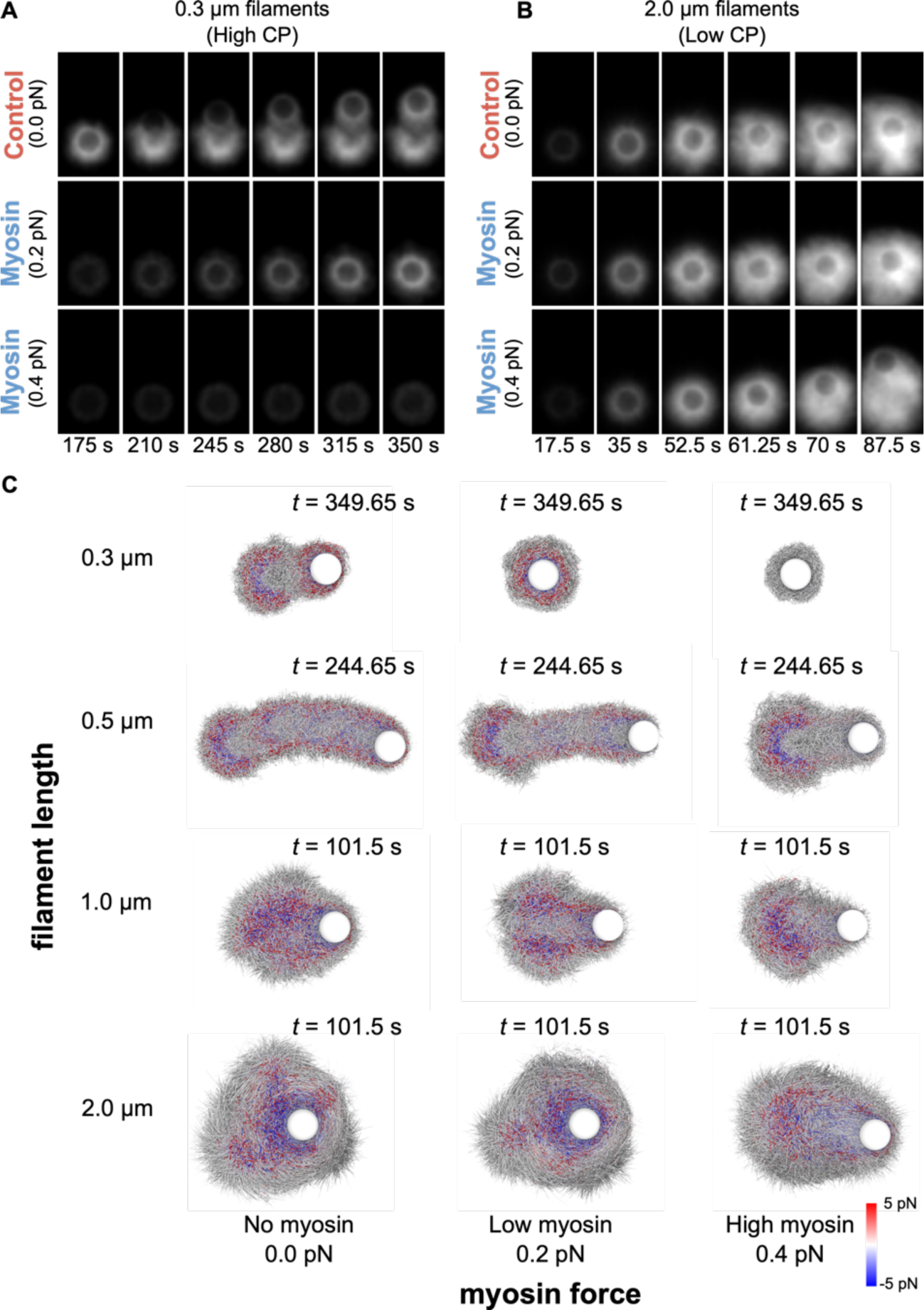
Simulated comet tail structure and tensile/compression force distributions as a function of branch length and myosin force *F*myo. (**A**) Simulated epifluorescence images as in Figure 7D, but with filaments grow to a length of 0.3 μm before capping. The simulations can be compared to high CP concentration experiments of Figure 2A where a comet tail fails to form for myosin-beads. (**B**) Same as in panel A, but for filament length of 2.0 μm. The simulations mimic the low CP concentration experiments of Figure 2B where myosin was found to promote symmetry breaking. (**C**) State diagram for actin comet tails as a function of filament length and myosin force. Images show a cut through the center of the comet tail. Color scale indicates filament tension (red: tensile; blue: compressive).

**Figure S6.**
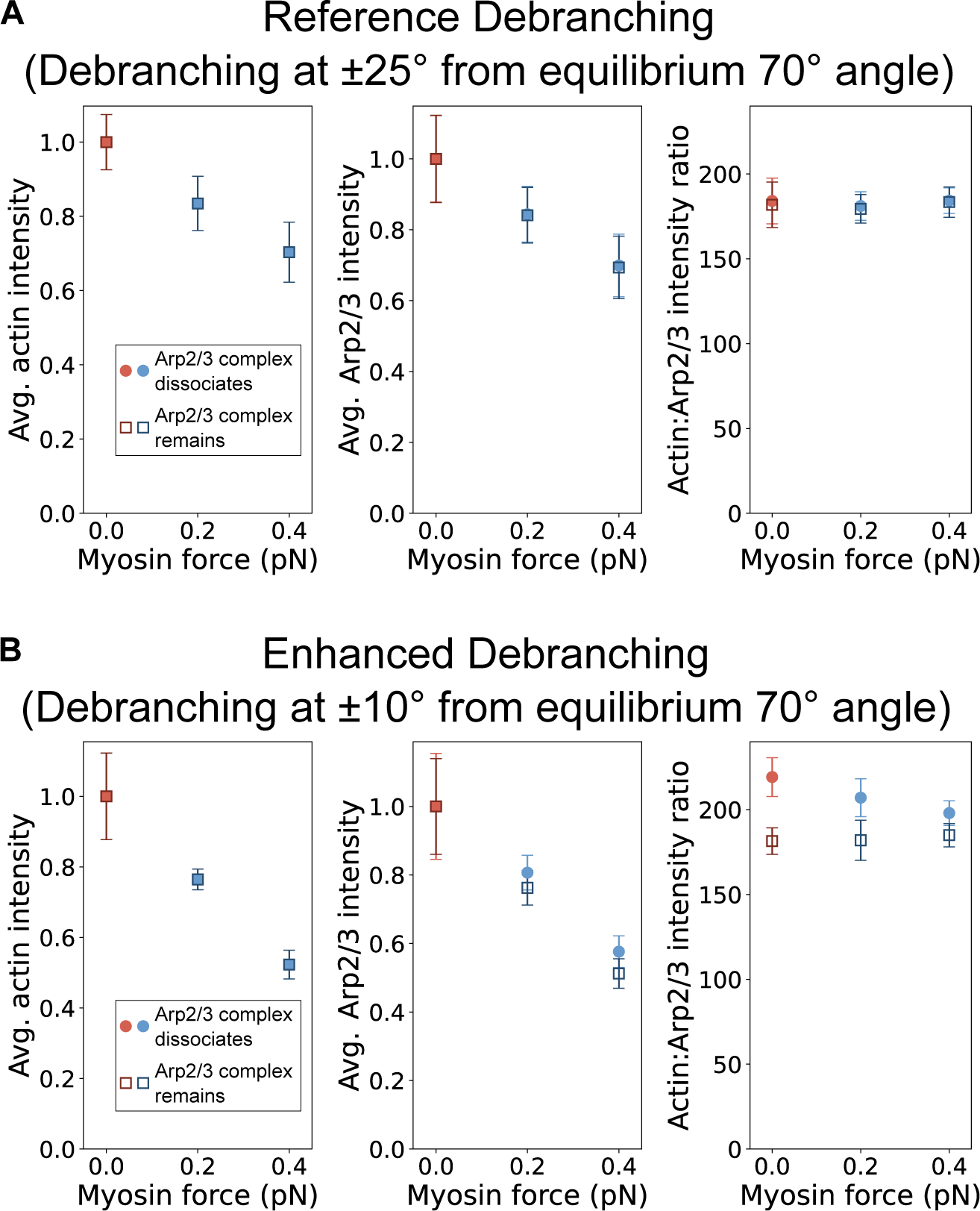
Simulated actin intensity, Arp2/3 intensity and Actin:Arp2/3 ratio across the entire actin comet tail for intermediate length condition (intermediate CP concentration) under different debranching thresholds. (**A**) Actin:Arp2/3 complex ratio (calibrated to number of actin monomers per Arp2/3 complex), average actin intensity, and average Arp2/3 complex intensity from simulated epifluorescent images of comet tails formed at reference debranching conditions with filament length 0.5 μm (≈187 subunits). Solid circles indicate averages calculated where the Arp2/3 complex is removed upon debranching; empty squares indicate averages calculated where the Arp2/3 complex remains attached to the daughter filament upon debranching. (**B**) Same as in panel A except under enhanced debranching condition (debranching occurring at ±10° away from 70° equilibrium branch instead of ±25° as in the reference case in A). The ratio varies with myosin force when Arp2/3 complex dissociates after debranching, but in the opposite manner as compared to the experimental results in Figure 4D.

**Figure S7.**
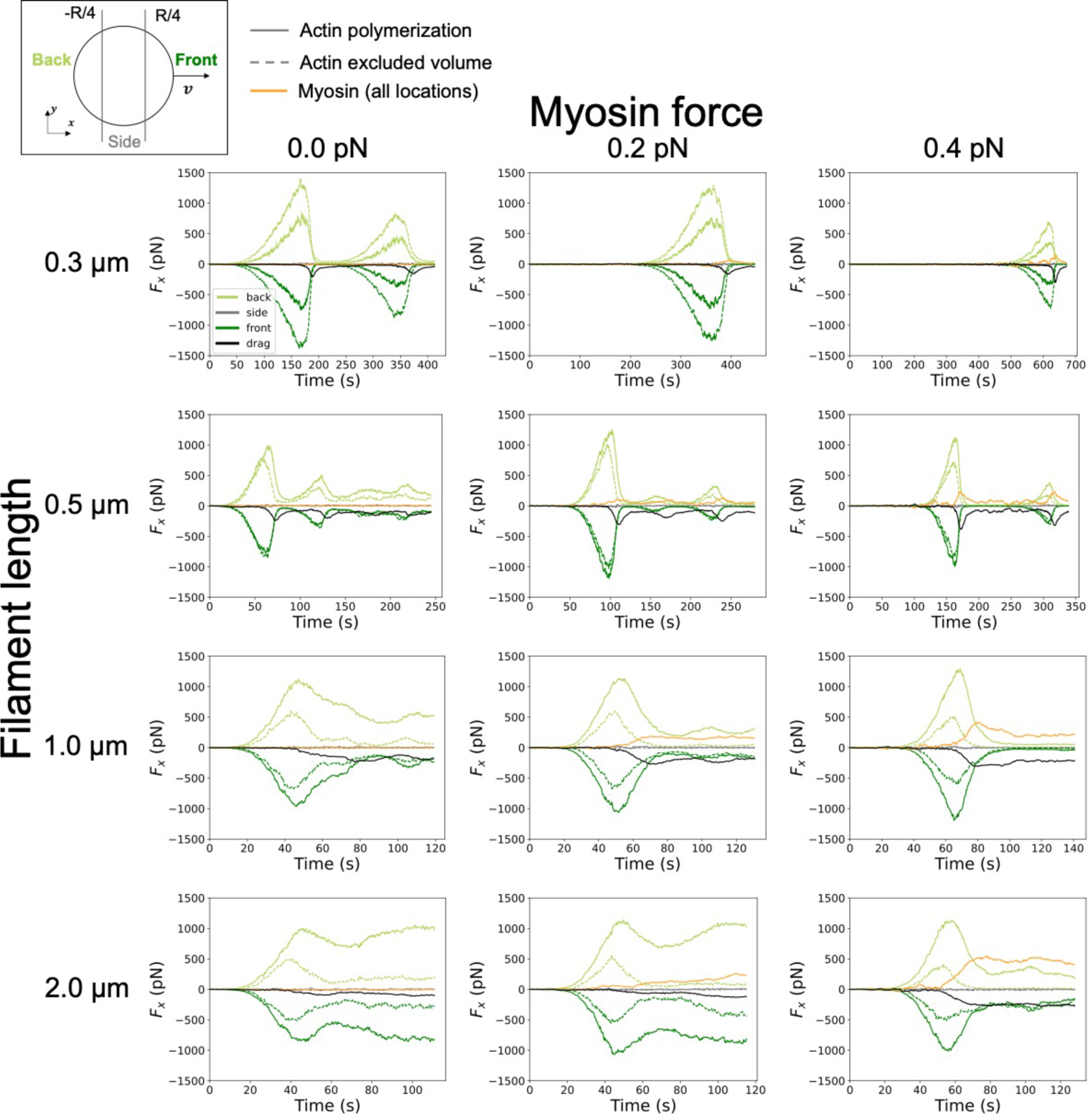
Force acting on beads during comet tail elongation. Subdivision of forces acting on beads along the direction of bead propulsion due to actin polymerization (light green, gray, dark green continuous; calculated as excluded volume interactions between barbed end segment and bead), actin excluded volume (light green, gray, dark green dashed; calculated as excluded volume interactions on the bead from filament segments other than barbed ends), and myosin forces on the bead (orange line). The bead drag force that balances the sum of these forces is shown in black. Actin forces are subdivided into whether they are located behind the bead (light green), at the side of the bead (gray), or in front of the bead (dark green).

**Figure S8.**
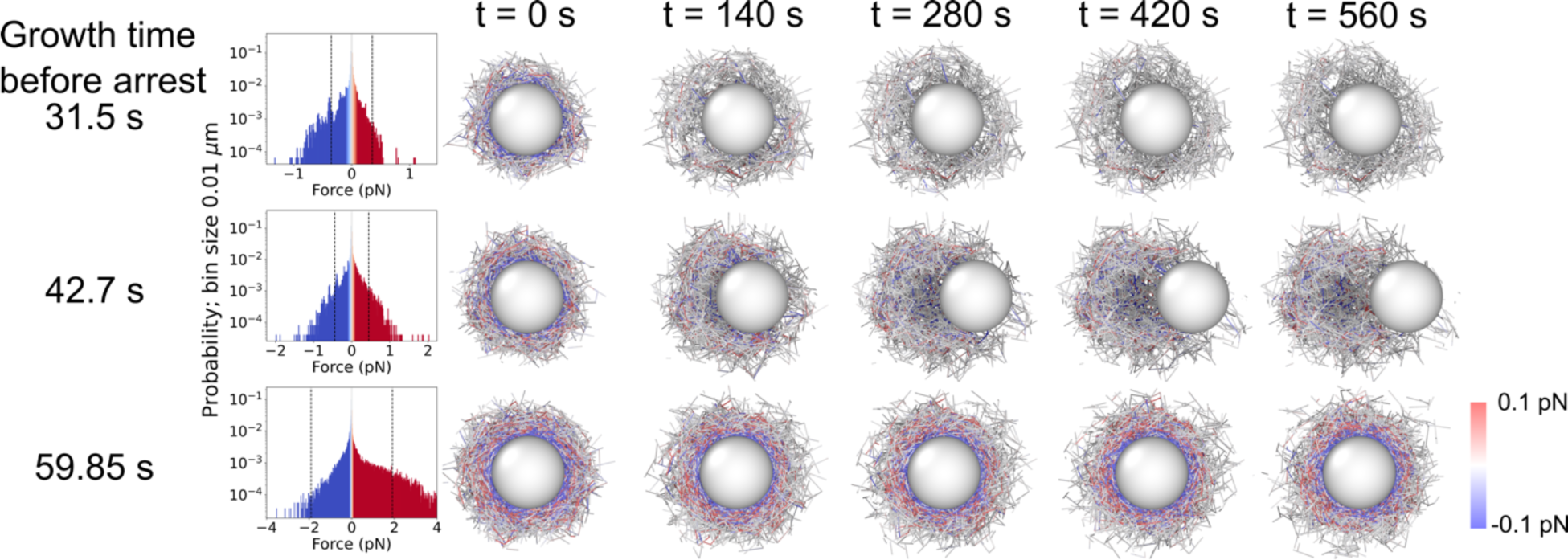
Simulated expulsion of bead after halting actin polymerization/branching. Simulation snapshots at intermediate filament length (0.5 μm) (intermediate CP concentration) show a cut through the bead center during the evolution of the actin network after halting actin polymerization and branching. Expulsion of the bead only occurs at intermediate cloud thickness (Growth time = 42.7 s, middle row and Fig. 7J). Thinner shells fall apart (Growth time = 31.5 s), while thicker shells cannot be broken by 0.2 pN myosin alone (Growth time = 59.85 s). Network evolved in the presence of the same myosin force (0.2 pN). Histograms show filament tension distribution at t = 0 s (time of growth arrest); dashed lines indicate region containing 95% of data, centered at zero.

**Table S1.** Table of simulation constants.

| Parameter | Description | Value | Justification |
| --- | --- | --- | --- |
| $R$ | Radius of nucleating bead | $0.75\ \mu\text{m}$ | Smaller but comparable to experiment, for numerical efficiency |
| $\Delta t$ | Simulation timestep | $3.5 \times 10^{-5}\ \text{s}$ | Allows for numerical stability of excluded volume interactions |
| $\zeta$ | Friction coefficient for actin segment | $0.377\ \text{pN s } \mu\text{m}^{-1}$ | Sufficiently high for numerical stability |
| $\zeta^{nucl\ bead}$ | Friction coefficient for nucleating bead | $2970\ \text{pN s } \mu\text{m}^{-1}$ | Sufficiently high for bead to not eject bare from actin shell |
| $l_p$ | Actin filament persistence length | $17.0\ \mu\text{m}$ | |
| $k^{actin}$ | Actin filament segment spring constant | $1000\ \text{pN } \mu\text{m}^{-1}$ | As large as allowed for numerical stability (62) |
| $l_0$ | Actin segment spring equilibrium distance | $0.1\ \mu\text{m}$ | Length of filament segment |
| $l_{00}$ | Starting filament length | $0.035\ \mu\text{m}$ | Numerical stability |
| $l^{branch}$ | Bond length between mother point element and pointed end of daughter filament | $0.02\ \mu\text{m}$ | Numerical stability |
| $d^{branch}$ | Range of branching nucleation | $0.1\ \mu\text{m}$ | Of order $l_0$ |
| $\epsilon_{angle}$ | Spring constant for filament branch | $2\ \text{pN } \mu\text{m}$ | Maintains angle between $60^\circ$-$80^\circ$ with thermal fluctuations (62) |
| $d^{excluded}$ | Excluded volume interaction range between filaments | $0.02\ \mu\text{m}$ | Larger than 7 nm filament diameter for increased numerical stability |
| $k^{excluded}$ | Excluded volume interaction constant | $1690\ \text{pN } \mu\text{m}^{-1}$ | Prior computational model (72) |
| $d^{myo}$ | Range of myosin force | $0.025\ \mu\text{m}$ | Comparable to size of myosin 1d |
| $F^{myo}$ | Magnitude of myosin force acting on actin filaments within range. | $0.0 - 0.4\ \text{pN}$ | Varied |
| $r_0^{pol}$ | Rate of polymerization of uncapped barbed ends | $40\ \text{sub s}^{-1}$ | $4\ \mu\text{M}$ actin solution |
| $r_0^{branch}$ | Rate of branching per filament point element | $0.15 \text{ s}^{-1}$ | Tuned to approximate timescale of symmetry breaking and comet speed (intermediate CP) |
| $r_0^{denovo}$ | Rate of spontaneous introduction of new filaments | $0.125 \text{ s}^{-1}$ | Tuned to allow actin shell on timescales comparable to experiment |
| $F^{frag}$ | Threshold force for tension fragmentation of filaments | 50 pN | Allows symmetry breaking |
| $F^{debranch}$ | Threshold force for tension debranching of branches | 30 pN | Of order $F^{frag}$ |
| $\Delta\theta^{debranch}$ | Threshold angle deviation from $70^\circ$ for debranching | $25^\circ$ | Debranching by bending and pulling (72) |

**Movie S1.** Myosin-bead (red) processively moving along a (green) single-actin-filament track.

**Movie S2.** Movie of control-beads and myosin-beads (0.43:1 #myosin/NPFs) acquired in the same imaging field in the presence of 200 nM CP showing the inability of myosin-beads to form a comet tail. Scale bar 5 µm.

**Movie S3.** Movie of control-beads and myosin-beads (0.43:1 #myosin/NPFs) acquired in the same field in the presence of 25 nM CP showing fracturing of the actin shell and comet tail growth from a myosin-bead but not the control-bead. Scale bar 5 µm.

**Movie S4.** Movie of control-beads and myosin-beads (0.43:1 #myosin/NPFs) growing comet tails in the presence of 50 nM CP. The actin network growing from the myosin-bead is less dense but has a similar tail length as the control. Scale bar 5 µm.

**Movie S5.** Movie of actin assembly around control-beads and rigor myosin-beads (0.43:1 #myosin/NPFs) at 50 nM CP. Rigor myosin heavily delayed the growth of actin comet tails. Scale bar 5 µm.

**Movie S6.** Movie of actin assembly around control- and HaloABD-bead at 50 nM CP. HaloABD of *α*-actinin heavily delayed the growth of actin comet tail as rigor myosin. Scale bar 5 µm.

**Movie S7.** Movie of myosin-beads (0.43:1 #myosin/NPFs) break actin shell apart acquired 11 min after actin polymerization was arrested by Latrunculin B and CK-666 in the absence of phalloidin. Scale bar 5 µm.

**Movie S8.** Movie of myosin-beads (0.43:1 #myosin/NPFs) ejected out of actin shell acquired 24 min after actin polymerization was arrested by Latrunculin B and CK-666 n the absence of phalloidin. Scale bar 5 µm.

**Movie S9** Simulated symmetry breaking and comet growth for intermediate length (0.5 μm) filaments, without myosin forces (Fig. 7B). Top video shows a cutaway (filament point elements with position z > 0 are not shown) while bottom video shows all filaments. Bead is semi-transparent.

**Movie S10.** Movie of simulated actin comet tail formation for short filaments (0.3 μm, High CP),_as a function of myosin force (Fig. S5A).

**Movie S11.** Movie of simulated actin comet tail formation for long filaments (2.0 μm, Low CP), as a function of myosin force (Fig. S5B).

**Movie S12.** Movie of simulated actin comet tail formation for intermediate filaments (0.5 μm, Intermediate CP), as a function of myosin force (Fig. 7D).

**Movie S13.** Movie of simulated symmetry breaking and bead ejection by myosin forces following arrest of actin polymerization and branching at 42.7 s (Fig 7I). Myosin force is 0.2 pN throughout the simulation and filament length is 0.5 μm.

## References

1. L. Blanchoin, R. Boujemaa-Paterski, C. Sykes, J. Plastino, Actin dynamics, architecture, and mechanics in cell motility. Physiol Rev 94, 235–263 (2014).

2. K. Rottner, J. Faix, S. Bogdan, S. Linder, E. Kerkhoff, Actin assembly mechanisms at a glance. Journal of Cell Science 130, 3427–3435 (2017).

3. T. Svitkina, The Actin Cytoskeleton and Actin-Based Motility. Cold Spring Harb Perspect Biol 10, (2018).

4. V. Papalazarou, L. M. Machesky, The cell pushes back: The Arp2/3 complex is a key orchestrator of cellular responses to environmental forces. Curr Opin Cell Biol 68, 37–44 (2021).

5. A. M. Gautreau, F. E. Fregoso, G. Simanov, R. Dominguez, Nucleation, stabilization, and disassembly of branched actin networks. Trends Cell Biol 32, 421–432 (2022).

6. M. Krendel, N. C. Gauthier, Building the phagocytic cup on an actin scaffold. Curr Opin Cell Biol 77, 102112 (2022).

7. M. Jin et al., Branched actin networks are organized for asymmetric force production during clathrin-mediated endocytosis in mammalian cells. Nat Commun 13, 3578 (2022).

8. D. N. Clarke, A. C. Martin, Actin-based force generation and cell adhesion in tissue morphogenesis. Curr Biol 31, R667–R680 (2021).

9. T. P. Loisel, R. Boujemaa, D. Pantaloni, M. F. Carlier, Reconstitution of actin-based motility of Listeria and Shigella using pure proteins. Nature 401, 613–616 (1999).

10. F. Nakamura, E. Osborn, P. A. Janmey, T. P. Stossel, Comparison of filamin A-induced cross-linking and Arp2/3 complex-mediated branching on the mechanics of actin filaments. J Biol Chem 277, 9148–9154 (2002).

11. O. Chaudhuri, S. H. Parekh, D. A. Fletcher, Reversible stress softening of actin networks. Nature 445, 295–298 (2007).

12. O. Akin, R. D. Mullins, Capping protein increases the rate of actin-based motility by promoting filament nucleation by the Arp2/3 complex. Cell 133, 841–851 (2008).

13. J. Stricker, T. Falzone, M. L. Gardel, Mechanics of the F-actin cytoskeleton. J Biomech 43, 9–14 (2010).

14. A. Kawska et al., How actin network dynamics control the onset of actin-based motility. Proc Natl Acad Sci U S A 109, 14440–14445 (2012).

15. P. Bieling et al., Force Feedback Controls Motor Activity and Mechanical Properties of Self-Assembling Branched Actin Networks. Cell 164, 115–127 (2016).

16. J. Mueller et al., Load Adaptation of Lamellipodial Actin Networks. Cell 171, 188–200 e116 (2017).

17. R. D. Mullins, P. Bieling, D. A. Fletcher, From solution to surface to filament: actin flux into branched networks. Biophys Rev 10, 1537–1551 (2018).

18. J. Funk, F. Merino, M. Schaks, K. Rottner, S. Raunser, P. Bieling, A barbed end interference mechanism reveals how capping protein promotes nucleation in branched actin networks. Nature Communications 12, (2021).

19. T. D. Li, P. Bieling, J. Weichsel, R. D. Mullins, D. A. Fletcher, The molecular mechanism of load adaptation by branched actin networks. Elife 11, (2022).

20. T. D. Pollard, E. D. Korn, Acanthamoeba Myosin. Journal of Biological Chemistry 248, 4682–4690 (1973).

21. B. B. McIntosh, E. M. Ostap, Myosin-I molecular motors at a glance. J Cell Sci 129, 2689–2695 (2016).

22. C. G. Almeida, A. Yamada, D. Tenza, D. Louvard, G. Raposo, E. Coudrier, Myosin 1b promotes the formation of post-Golgi carriers by regulating actin assembly and membrane remodelling at the trans-Golgi network. Nat Cell Biol 13, 779–789 (2011).

23. R. T. A. Pedersen, D. G. Drubin, Type I myosins anchor actin assembly to the plasma membrane during clathrin-mediated endocytosis. J Cell Biol 218, 1138–1147 (2019).

24. H. E. Manenschijn, A. Picco, M. Mund, A. S. Rivier-Cordey, J. Ries, M. Kaksonen, Type-I myosins promote actin polymerization to drive membrane bending in endocytosis. Elife 8, (2019).

25. A. Capmany et al., MYO1C stabilizes actin and facilitates the arrival of transport carriers at the Golgi complex. J Cell Sci 132, (2019).

26. S. R. Barger et al., Membrane-cytoskeletal crosstalk mediated by myosin-I regulates adhesion turnover during phagocytosis. Nat Commun 10, 1249 (2019).

27. M. J. Greenberg, E. M. Ostap, Regulation and control of myosin-I by the motor and light chain-binding domains. Trends Cell Biol 23, 81–89 (2013).

28. R. Boujemaa-Paterski, R. Galland, C. Suarez, C. Guerin, M. Thery, L. Blanchoin, Directed actin assembly and motility. Methods Enzymol 540, 283–300 (2014).

29. V. Noireaux et al., Growing an actin gel on spherical surfaces. Biophys J 78, 1643–1654 (2000).

30. J. H. Lewis, T. Lin, D. E. Hokanson, E. M. Ostap, Temperature dependence of nucleotide association and kinetic characterization of myo1b. Biochemistry 45, 11589–11597 (2006).

31. F. A. Baez-Cruz, E. M. Ostap, Drosophila class-I myosins that can impact left-right asymmetry have distinct ATPase kinetics. J Biol Chem 299, 104961 (2023).

32. L. A. Cameron, M. J. Footer, A. van Oudenaarden, J. A. Theriot, Motility of ActA protein-coated microspheres driven by actin polymerization. Proc Natl Acad Sci U S A 96, 4908–4913 (1999).

33. A. Bernheim-Groswasser, S. Wiesner, R. M. Golsteyn, M. F. Carlier, C. Sykes, The dynamics of actin-based motility depend on surface parameters. Nature 417, 308–311 (2002).

34. J. van der Gucht, E. Paluch, J. Plastino, C. Sykes, Stress release drives symmetry breaking for actin-based movement. Proc Natl Acad Sci U S A 102, 7847–7852 (2005).

35. V. Achard et al., A “primer”-based mechanism underlies branched actin filament network formation and motility. Curr Biol 20, 423–428 (2010).

36. M. J. Dayel, O. Akin, M. Landeryou, V. Risca, A. Mogilner, R. D. Mullins, In silico reconstitution of actin-based symmetry breaking and motility. PLoS Biol 7, e1000201 (2009).

37. D. H. Wachsstock, W. H. Schwarz, T. D. Pollard, Cross-linker dynamics determine the mechanical properties of actin gels. Biophys J 66, 801–809 (1994).

38. P. A. Kuhlman, J. Ellis, D. R. Critchley, C. R. Bagshaw, The kinetics of the interaction between the actin-binding domain of alpha-actinin and F-actin. FEBS Lett 339, 297–301 (1994).

39. J. H. Lewis, M. J. Greenberg, J. M. Laakso, H. Shuman, E. M. Ostap, Calcium regulation of myosin-I tension sensing. Biophys J 102, 2799–2807 (2012).

40. S. Manceva, T. Lin, H. Pham, J. H. Lewis, Y. E. Goldman, E. M. Ostap, Calcium regulation of calmodulin binding to and dissociation from the myo1c regulatory domain. Biochemistry 46, 11718–11726 (2007).

41. B. J. Nolen et al., Characterization of two classes of small molecule inhibitors of Arp2/3 complex. Nature 460, 1031–1034 (2009).

42. P. Bieling et al., WH2 and proline-rich domains of WASP-family proteins collaborate to accelerate actin filament elongation. EMBO J 37, 102–121 (2018).

43. N. G. Pandit et al., Force and phosphate release from Arp2/3 complex promote dissociation of actin filament branches. Proc Natl Acad Sci U S A 117, 13519–13528 (2020).

44. J. Pernier et al., Myosin 1b flattens and prunes branched actin filaments. J Cell Sci 133, (2020).

45. R. T. A. Pedersen, A. Snoberger, S. Pyrpassopoulos, D. Safer, D. G. Drubin, E. M. Ostap, Endocytic myosin-1 is a force-insensitive, power-generating motor. J Cell Biol 222, (2023).

46. F. S. Wang, C. W. Liu, T. J. Diefenbach, D. G. Jay, Modeling the role of myosin 1c in neuronal growth cone turning. Biophys J 85, 3319–3328 (2003).

47. J. L. Maravillas-Montero, P. G. Gillespie, G. Patino-Lopez, S. Shaw, L. Santos-Argumedo, Myosin 1c participates in B cell cytoskeleton rearrangements, is recruited to the immunologic synapse, and contributes to antigen presentation. J Immunol 187, 3053–3063 (2011).

48. A. M. Sokac, C. Schietroma, C. B. Gundersen, W. M. Bement, Myosin-1c couples assembling actin to membranes to drive compensatory endocytosis. Dev Cell 11, 629–640 (2006).

49. J. Cheng, A. Grassart, D. G. Drubin, Myosin 1E coordinates actin assembly and cargo trafficking during clathrin-mediated endocytosis. Mol Biol Cell 23, 2891–2904 (2012).

50. M. Krendel, E. K. Osterweil, M. S. Mooseker, Myosin 1E interacts with synaptojanin-1 and dynamin and is involved in endocytosis. FEBS Lett 581, 644–650 (2007).

51. J. L. Ouderkirk, M. Krendel, Myosin 1e is a component of the invadosome core that contributes to regulation of invadosome dynamics. Exp Cell Res 322, 265–276 (2014).

52. M. E. Garone, S. E. Chase, C. Zhang, M. Krendel, Myosin 1e deficiency affects migration of 4T1 breast cancer cells. Cytoskeleton (Hoboken), (2023).

53. Y. Sun, A. C. Martin, D. G. Drubin, Endocytic internalization in budding yeast requires coordinated actin nucleation and myosin motor activity. Dev Cell 11, 33–46 (2006).

54. J. M. Laakso, J. H. Lewis, H. Shuman, E. M. Ostap, Myosin I can act as a molecular force sensor. Science 321, 133–136 (2008).

55. J. A. Spudich, S. Watt, The regulation of rabbit skeletal muscle contraction. I. Biochemical studies of the interaction of the tropomyosin-troponin complex with actin and the proteolytic fragments of myosin. J Biol Chem 246, 4866–4871 (1971).

56. D. R. Kellogg, T. J. Mitchison, B. M. Alberts, Behaviour of microtubules and actin filaments in living Drosophila embryos. Development 103, 675–686 (1988).

57. G. Lebreton et al., Molecular to organismal chirality is induced by the conserved myosin 1D. Science 362, 949–952 (2018).

58. M. Boczkowska, G. Rebowski, M. V. Petoukhov, D. B. Hayes, D. I. Svergun, R. Dominguez, X-ray scattering study of activated Arp2/3 complex with bound actin-WCA. Structure 16, 695–704 (2008).

59. A. Zimmet, T. Van Eeuwen, M. Boczkowska, G. Rebowski, K. Murakami, R. Dominguez, Cryo-EM structure of NPF-bound human Arp2/3 complex and activation mechanism. Sci Adv 6, (2020).

60. J. N. Rao, Y. Madasu, R. Dominguez, Mechanism of actin filament pointed-end capping by tropomodulin. Science 345, 463–467 (2014).

61. M. J. Greenberg, H. Shuman, E. M. Ostap, Measuring the Kinetic and Mechanical Properties of Non-processive Myosins Using Optical Tweezers. Methods Mol Biol 1486, 483–509 (2017).

62. D. M. Rutkowski, D. Vavylonis, Discrete mechanical model of lamellipodial actin network implements molecular clutch mechanism and generates arcs and microspikes. PLoS Comput Biol 17, e1009506 (2021).

63. A. E. Carlsson, Growth of branched actin networks against obstacles. Biophys J 81, 1907–1923 (2001).

64. J. B. Alberts, G. M. Odell, In silico reconstitution of Listeria propulsion exhibits nano-saltation. PLoS Biol 2, e412 (2004).

65. D. Holz, D. Vavylonis, Building a dendritic actin filament network branch by branch: models of filament orientation pattern and force generation in lamellipodia. Biophys Rev 10, 1577–1585 (2018).

66. K. Sekimoto, J. Prost, F. Julicher, H. Boukellal, A. Bernheim-Grosswasser, Role of tensile stress in actin gels and a symmetry-breaking instability. Eur Phys J E Soft Matter 13, 247–259 (2004).

67. K. John, P. Peyla, K. Kassner, J. Prost, C. Misbah, Nonlinear study of symmetry breaking in actin gels: implications for cellular motility. Phys Rev Lett 100, 068101 (2008).

68. F. Gerbal, P. Chaikin, Y. Rabin, J. Prost, An elastic analysis of Listeria monocytogenes propulsion. Biophys J 79, 2259–2275 (2000).

69. J. Zhu, A. Mogilner, Mesoscopic model of actin-based propulsion. PLoS Comput Biol 8, e1002764 (2012).

70. N. J. Burroughs, D. Marenduzzo, Nonequilibrium-driven motion in actin networks: comet tails and moving beads. Phys Rev Lett 98, 238302 (2007).

71. K. C. Lee, A. J. Liu, New Proposed Mechanism of Actin-Polymerization-Driven Motility. Biophysical Journal 95, 4529–4539 (2008).

72. T. Kim, W. Hwang, H. Lee, R. D. Kamm, Computational analysis of viscoelastic properties of crosslinked actin networks. PLoS Comput Biol 5, e1000439 (2009).

73. C. S. Peskin, G. M. Odell, G. F. Oster, Cellular motions and thermal fluctuations: the Brownian ratchet. Biophys J 65, 316–324 (1993).

74. A. Mogilner, G. Oster, Cell motility driven by actin polymerization. Biophys J 71, 3030–3045 (1996).

75. Y. Tsuda, H. Yasutake, A. Ishijima, T. Yanagida, Torsional rigidity of single actin filaments and actin-actin bond breaking force under torsion measured directly by in vitro micromanipulation. Proc Natl Acad Sci U S A 93, 12937–12942 (1996).

